# Spine apparatus modulates Ca^2+^ in spines through spatial localization of sources and sinks

**DOI:** 10.1101/2023.09.22.558941

**Authors:** M. Herńandez Mesa, G. C. Garcia, F. J. Hoerndli, K. J. McCabe, P. Rangamani

## Abstract

Dendritic spines are small protrusions on dendrites in neurons and serve as sites of postsynaptic activity. Some of these spines contain smooth endoplasmic reticulum (SER), and sometimes an even further specialized SER known as the spine apparatus (SA). In this work, we developed a stochastic spatial model to investigate the role of the SER and the SA in modulating Ca^2+^ dynamics. Using this model, we investigated how ryanodine receptor (RyR) localization, spine membrane geometry, and SER geometry can impact Ca^2+^ transients in the spine and in the dendrite. Our simulations found that RyR opening is dependent on where it is localized in the SER and on the SER geometry. In order to maximize Ca^2+^ in the dendrites (for activating clusters of spines and spine-spine communication), a laminar SA was favorable with RyRs localized in the neck region, closer to the dendrite. We also found that the presence of the SER without the laminar structure, coupled with RyR localization at the head, leads to higher Ca^2+^ presence in the spine. These predictions serve as design principles for understanding how spines with an ER can regulate Ca^2+^ dynamics differently from spines without ER through a combination of geometry and receptor localization.

**Highlights:**

- RyR opening in dendritic spine ER is location dependent and spine geometry dependent.
- Ca^2+^ buffers and SERCA can buffer against runaway potentiation of spines even when CICR is activated.
- RyRs located towards the ER neck allow for more Ca^2+^ to reach the dendrites.
- RyRs located towards the spine head are favorable for increased Ca^2+^ in spines.

Figure 1:
Graphical abstract.
Factors governing the dynamics of Ca^2+^ in dendritic spines include plasma membrane geometry, RyR distribution and ER laminarity.

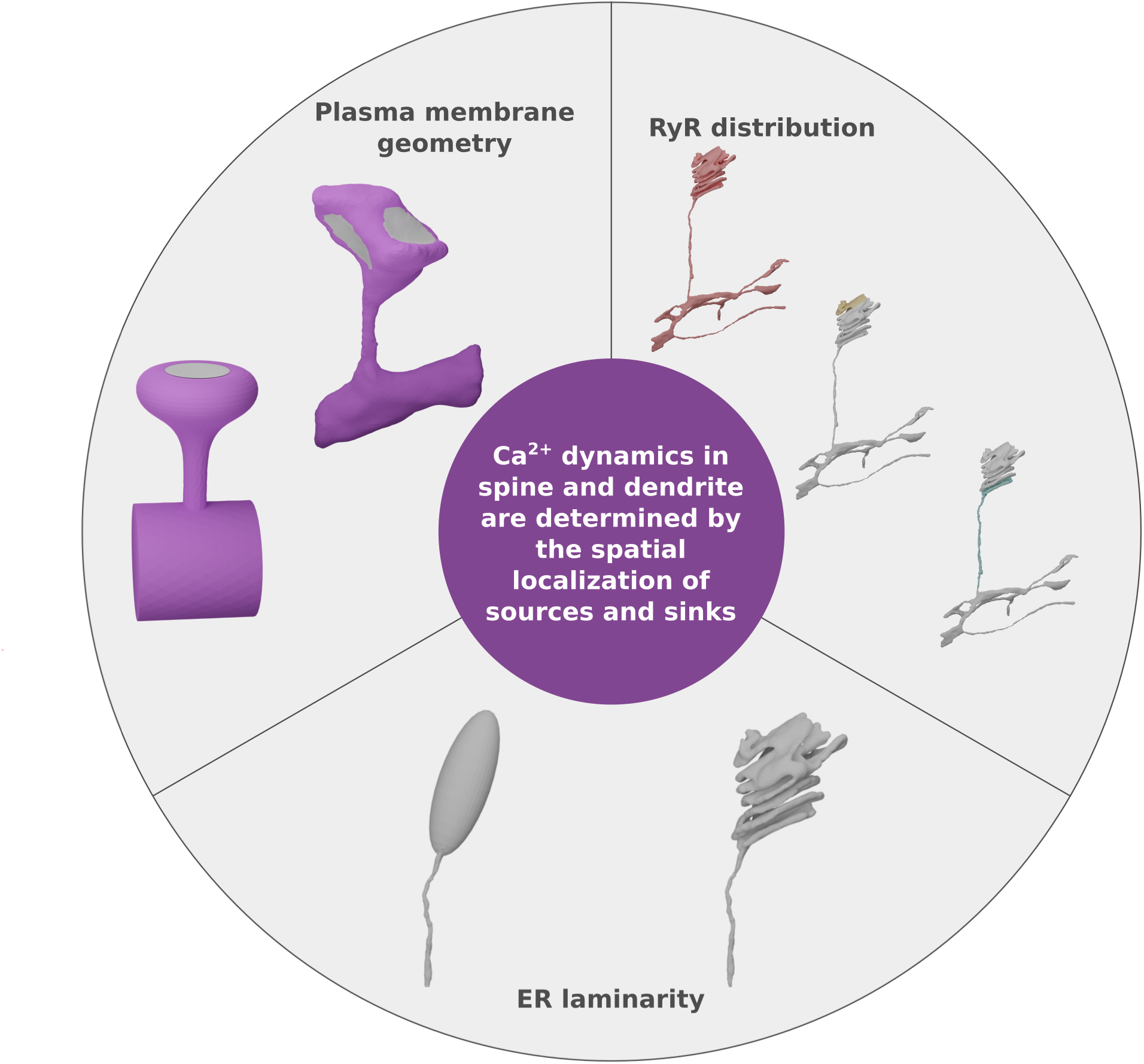

## 3 Introduction

Communication between two neurons occurs at the synapse, which consists of three parts: a presynaptic terminal on one neuron, the postsynaptic site of another neuron, and the synaptic cleft between them. Dendritic spines are small protrusions (0.04 to 0.29 µm^3^ (*1*, *2*)) located along the dendrite of the neuron receiving the signal (postsynaptic sites). The biochemical machinery initiated by the release of neurotransmitters activates multiple biochemical reactions within the spines (*3*), which play an important role in learning and memory formation (*4*). Dendritic spines are highly dynamic structures, capable of changing their size and shape in response to synaptic activity (*5* –*7*), and these alterations may affect learning and memory formation (*8*, *9*). Spines also come in different shapes and sizes and spine geometry is thought to impact synaptic plasticity (*1*, *4*, *10* –*12*). Ca^2+^ dynamics are a key determinant of learning and memory formation in spines as they regulate several mechanisms such as activation of Ca^2+^ sensitive proteins as Ca^2+^/calmodulin-dependent kinase type II (CaMKII), synaptic plasticity, and synapse-to-nucleus communication (*13*, *14*).

Synaptic Ca^2+^ signals are generated upon membrane depolarization as a response to release of glutamate from the presynaptic terminal (*14*). This allows for the activation of N-methyl-D-aspartate receptors (NMDAR) resulting in an influx of Ca^2+^. The voltage sensitive Ca^2+^ channels (VSCC) located on the plasma membrane of dendritic spines are an additional source for cytosolic Ca^2+^. In response to this Ca^2+^ influx, plasma membrane Ca^2+^-ATPase (PMCA) pumps and Na^+^-Ca^2+^ exchangers (NCX) remove the cytosolic Ca^2+^ out of the spine (Figure 2A). Intracellular Ca^2+^ stores can also modulate the levels of cytosolic Ca^2+^ concentrations, for example the endoplasmic reticulum (ER), which is the largest intracellular Ca^2+^ storage organelle in eukaryotic cells (*15*) (Figure 2A). Additionally, a subset of dendritic spines (around 48% of adult spines) contains smooth endoplasmic reticulum (SER) (*2*). A specialized form of the SER, called the spine apparatus (SA) (*16*, *17*), is identified by its characteristic laminar structure of stacked SA discs (*17*) and is found in about 10-20 % of adult spines (*2*). Spacek and Harris showed that the presence of SA is more prevalent in mature mushroom-shaped spines (*2*). The actin-associated protein synaptopodin is necessary for SA formation since it localizes to the F-actin matrix in between the SA discs (*18*). Due to its capacity for storing Ca^2+^, the SA has been suggested to play an important role in synaptic plasticity and Ca^2+^ signalling (*16*, *18*). Ca^2+^ is stored in the SA *via* the sarco/endoplasmic reticulum Ca^2+^-ATPase (SERCA) pump, while Ca^2+^ is released from the SA through the inositol 1,4,5 triphosphate receptor (IP_3_R) and the ryanodine receptor (RyR). Cytosolic Ca^2+^ ions activate RyRs and IP_3_Rs which allows for further Ca^2+^ release into the cytosol, a process known as Ca^2+^-induced Ca^2+^ release (CICR).

**Figure 2:**
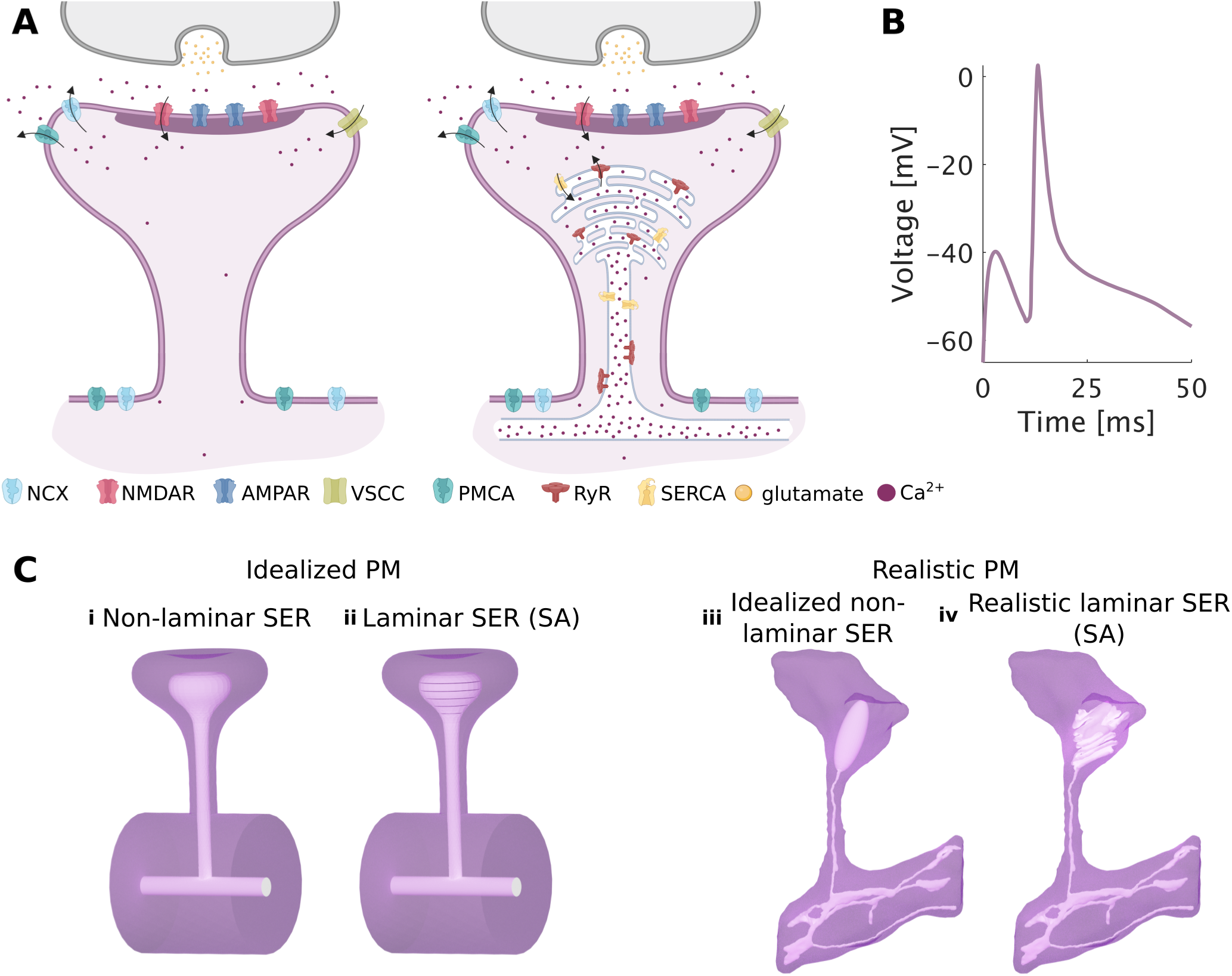
Overview of the model and geometries used to investigate the role of the SER and spine apparatus in regulating Ca^2+^ dynamics. **A**: Schematic of the molecular components, currents, and receptors included in this model. On the left, the spine apparatus is not included. On the right, a spine apparatus with RyRs and SERCA is included in the spine. The shaded part at the spines represents the postsynaptic density (PSD). This panel was generated using *Biorender.com*. **B**: The stimulus is modeled as glutamate release coupled with a simultaneous change in membrane potential characterized by an excitatory postsynaptic potential (EPSP) and a back-propagating action potential (bAP) as described in (*4*) and (*19*). **C**: The four different geometries used in this work consider both an idealized (*10*) and a realistic plasma membrane generated from experimental images (*20*), along with a non-laminar SER and a laminar SER representing the SA.

The localization of these receptors varies within the brain, with IP_3_Rs being more prevalent in dendritic shafts and RyRs more common in dendritic spines, as shown by Sharp *et al.* using immunohistochemistry (*21*, *22*). Although the effect of IP_3_Rs on plasticity has been extensively studied (*22* –*24*), the role of RyRs on synaptic plasticity remains to be fully explored. Korkotian and Segal have studied the role of the SA in Ca^2+^ dynamics using flashed photolysis of caged Ca^2+^ inside dendritic spines (*22*, *25*). They observed a second peak of cytosolic Ca^2+^ in spines with SA after flash photolysis of caged Ca^2+^ in dendritic spines; this second peak was not present in those spines lacking SA. When an RyR antagonist was used, the second peak was abolished in SA-containing spines (*22*, *25*). RyR localization within dendritic spines has been recently studied. Vlachos *et al.* showed experimentally the colocalization of the RyRs with synaptopodin (*26*) while Basnayake *et al.* (*27*) showed using super-resolution stimulated emission depletion imaging, in both cultured rat neurons and mouse hippocampal brain slices, that RyRs tend to be located on the lower part of the SA towards the base of the spine neck. However, the exact contribution of the SER, the SA and the localized pools of RyRs to synaptic plasticity and Ca^2+^ dynamics in intracellular signaling remains unclear (*28*).

Computational models can augment our understanding of the role of the SA and the RyRs on Ca^2+^ dynamics in dendritic spine communication. Breit *et al.* used a deterministic model to establish the importance of a SA with Ca^2+^ releasing receptors (RyRs and IP_3_Rs) in order for Ca^2+^ to reach the dendrites and thus allow spine-to-dendrite communication (*29*). Due to the small volume of spines, the use of stochastic models including particle diffusion instead of deterministic models has been deemed preferable (*19*, *30*, *31*). Therefore, Bell and Holst *et al.* used a stochastic model to study the effect of spine shape and SER morphology on synaptic weight (*4*). Bell *et al.* used a physiological stimulus based on glutamate release from a presynaptic terminal and membrane depolarization (*4*) to investigate how the volume-to-surface area of the spine modulates changes in synaptic strength on both spines without SER and on spines containing a passive SER. Separately, Basnayake *et al.* (*27*) also included stochastic simulations to show that more Ca^2+^ reaches the dendrite when the RyRs are located towards the lower part of the spine apparatus.

These studies highlight open questions about the impact of spine geometry, SER and SA geometry, and RyR localization on Ca^2+^ dynamics in spines and dendrites. For example, how does RyR localization on the SER affect Ca^2+^ dynamics? What configurations of SER geometry and RyR localization give rise to increased Ca^2+^ in the spine head? In the dendrite? To investigate the crosstalk between the geometric features listed above and Ca^2+^ dynamics, we developed a stochastic computational model of the main molecular components in spines with a focus on RyR-mediated CICR. In our model, we considered the role of RyR clustering inspired by work in cardiomyocytes (*32* –*34*) and the role of RyR localization based on experimental observations (*27*). Since the ER and the spine have unique shapes (*10*, *16*, *17*), we also investigated how the geometries of both the spine head and the ER (non-laminar and SA) could impact the RyR-driven Ca^2+^ dynamics using idealized and realistic geometries. Our simulations reveal complex rules for Ca^2+^ handling in dendritic spines and dendrites, foremost among which is that spine geometry and spine apparatus laminarity, in combination with receptor localization, govern RyR-mediated Ca^2+^ dynamics in spine heads and dendrites. This interplay is also affected by SERCA dynamics and Ca^2+^ buffering, which play important roles in modulating Ca^2+^ in spines and in dendrites. Taken together, these results provide novel insights into geometrical nuances and transport phenomena that ultimately affect downstream signal propagation in neurons.

## 4 Model development

In this work, we developed a stochastic spatial particle-based model that incorporates RyR dynamics into a previously described Ca^2+^ framework (*4*, *19*). In the following section we explain the details, methods, and assumptions taken with regard to model development. The receptors and channels considered in the model are depicted in Figure 2A. This section is divided as follows: first, we describe the equations used for the receptors at the plasma membrane. Then, we explain the equations in the cytosol and those used for describing the receptors used in the ER/SA. In the last two sections we describe the geometries used in this work, followed by the stimulus and further simulation details. An overview of all the equations used is given in Table 1 and in Section 4.3.1 for the RyR equations. The parameters used are summarized in Table 2. The model was implemented in the spatial, stochastic, particle-based framework MCell4 (*37*) in order to capture the stochasticity due to particle diffusion in small volumes such as spines. Since most of the equations were derived from Bell *et al.* (*4*), who developed their model using MCell3, we compared their results with our implementation of the model in the latest version of MCell to ensure that the results matched, see Figure S1. Additionally, an ODE version was implemented in MATLAB (*38*) to allow for the optimization of specific global parameters.

**Table 1:** Chemical reactions used in the model, with the exception of the the RyR equations, which are listed in Section 4.3.1.

| Reactions for NMDAR | Ref. | Reactions for PMCA | Ref. |
| --- | --- | --- | --- |
| $\text{NMDAR}_0 + \text{glut.} \xrightleftharpoons[k_{N,C1C0}]{k_{N,C0C1}} \text{NMDAR}_1$ | (35) | $\text{Ca}_{\text{cyto}}^{2+} + \text{PMCA} \xrightleftharpoons[k_{P2}]{k_{P1}} \text{PMCA}_1$ | (19) |
| $\text{NMDAR}_1 + \text{glut.} \xrightleftharpoons[k_{N,C2C1}]{k_{N,C1C2}} \text{NMDAR}_2$ | (35) | $\text{PMCA}_1 \xrightarrow{k_{P3}} \text{PMCA}$ | (19) |
| $\text{NMDAR}_2 \xrightleftharpoons[k_{DC2}]{k_{C2D}} \text{NMDAR}_D$ | (35) | $\text{PMCA} \xrightarrow{k_{P,\text{leak}}} \text{PMCA} + \text{Ca}_{\text{cyto}}^{2+}$ | (19) |
| $\text{NMDAR}_2 \xrightleftharpoons[k_{N,0C2}]{k_{N,C20}} \text{NMDAR}$ | (35) | Reactions for NCX | Ref. |
| $\text{NMDAR} \xrightarrow{k_{N,(V_m)}} \text{NMDAR} + \text{Ca}_{\text{cyto}}^{2+}$ | (35) | $\text{Ca}_{\text{cyto}}^{2+} + \text{NCX} \xrightleftharpoons[k_{N2}]{k_{N1}} \text{NCX}_1$ | (19) |
| $\text{NMDAR}_B \xrightleftharpoons[k_{B,(V_m)}]{k_{U,(V_m)}} \text{NMDAR}$ | (35) | $\text{NCX}_1 \xrightarrow{k_{N3}} \text{NCX}$ | (19) |
| $\text{NMDAR}_{2B} \xrightleftharpoons[k_{0C2b}]{k_{C20b}} \text{NMDAR}_B$ | (35) | $\text{NCX} \xrightarrow{k_{N,\text{leak}}} \text{NCX} + \text{Ca}_{\text{cyto}}^{2+}$ | (19) |
| $\text{NMDAR}_{2B} \xrightleftharpoons[k_{DC2}]{k_{C2D}} \text{NMDAR}_{DB}$ | (35) | Reactions for $\text{Ca}^{2+}$ buffers | Ref. |
| $\text{NMDAR}_{1B} + \text{glut.} \xrightleftharpoons[k_{N,C2C1}]{k_{N,C1C2}} \text{NMDAR}_{2B}$ | (35) | $\text{Ca}_{\text{cyto}}^{2+} + \text{B}_f \xrightleftharpoons[k_{Bf,\text{off}}]{k_{Bf,\text{on}}} \text{CaB}_f$ | (4) |
| $\text{NMDAR}_{0B} + \text{glut.} \xrightleftharpoons[k_{N,C1C0}]{k_{N,C0C1}} \text{NMDAR}_{1B}$ | (35) | $\text{Ca}_{\text{cyto}}^{2+} + \text{B}_m \xrightleftharpoons[k_{Bm,\text{off}}]{k_{Bm,\text{on}}} \text{CaB}_m$ | (4) |
| Reactions for AMPAR | Ref. | Reactions for $\text{Ca}^{2+}$ decay | Ref. |
| $\text{AMPAR}_0 + \text{glut.} \xrightleftharpoons[k_{A,C1C0}]{k_{A,C0C1}} \text{AMPAR}_1$ | (36) | $\text{Ca}_{\text{cyto}}^{2+} \xrightarrow{k_d} \emptyset$ | (4) |
| $\text{AMPAR}_1 + \text{glut.} \xrightleftharpoons[k_{A,C2C1}]{k_{A,C1C2}} \text{AMPAR}_2$ | (36) | Reactions for SERCA | Ref. |
| $\text{AMPAR}_2 \xrightleftharpoons[k_{A,C02}]{k_{A,C20}} \text{AMPAR}$ | (36) | $\text{SERCA}_x + \text{Ca}_{\text{cyto}}^{2+} \xrightleftharpoons[k_{x1x0}]{k_{x0x1}} \text{SERCA}_{x1}$ | (19) |
| $\text{AMPAR}_1 \xrightleftharpoons[k_{C3C1}]{k_{C1C3}} \text{AMPAR}_3$ | (36) | $\text{SERCA}_{x1} + \text{Ca}_{\text{cyto}}^{2+} \xrightleftharpoons[k_{x2x1}]{k_{x1x2}} \text{SERCA}_{x2}$ | (19) |
| $\text{AMPAR}_3 + \text{glut.e} \xrightleftharpoons[k_{C4C3}]{k_{C3C4}} \text{AMPAR}_4$ | (36) | $\text{SERCA}_{x2} \xrightleftharpoons[k_{y2x2}]{k_{x2y2}} \text{SERCA}_{y2}$ | (19) |
| $\text{AMPAR}_2 \xrightleftharpoons[k_{C4C2}]{k_{C2C4}} \text{AMPAR}_4$ | (36) | $\text{SERCA}_{y2} \xrightleftharpoons[k_{y1y2}]{k_{y2y1}} \text{SERCA}_{y1} + \text{Ca}_{\text{ER}}^{2+}$ | (19) |
| $\text{AMPAR}_4 \xrightleftharpoons[k_{C5C4}]{k_{C4C5}} \text{AMPAR}_5$ | (36) | $\text{SERCA}_{y1} \xrightleftharpoons[k_{y0y1}]{k_{y1y0}} \text{SERCA}_y + \text{Ca}_{\text{ER}}^{2+}$ | (19) |
| $\text{AMPAR}_5 \xrightleftharpoons[k_{0C5}]{k_{C50}} \text{AMPAR}$ | (36) | $\text{SERCA}_y \xrightleftharpoons[k_{x0y0}]{k_{y0x0}} \text{SERCA}_x$ | (19) |
| Reactions for VSCC | Ref. | $\text{Ca}_{\text{ER}}^{2+} \xrightarrow{k_{S,\text{leak}}} \text{Ca}_{\text{cyto}}^{2+}$ | (4) |
| $\text{VSCC}_0 \xrightleftharpoons[b_1(V_m)]{a_1(V_m)} \text{VSCC}_1$ | (19) | | |
| $\text{VSCC}_1 \xrightleftharpoons[b_2(V_m)]{a_2(V_m)} \text{VSCC}_2$ | (19) | | |
| $\text{VSCC}_2 \xrightleftharpoons[b_3(V_m)]{a_3(V_m)} \text{VSCC}_3$ | (19) | | |
| $\text{VSCC}_3 \xrightleftharpoons[b_4(V_m)]{a_4(V_m)} \text{VSCC}$ | (19) | | |
| $\text{VSCC} \xrightarrow{k_{Ca(V_m)}} \text{VSCC} + \text{Ca}_{\text{cyto}}^{2+}$ | (19) | | |

**Table 2:**
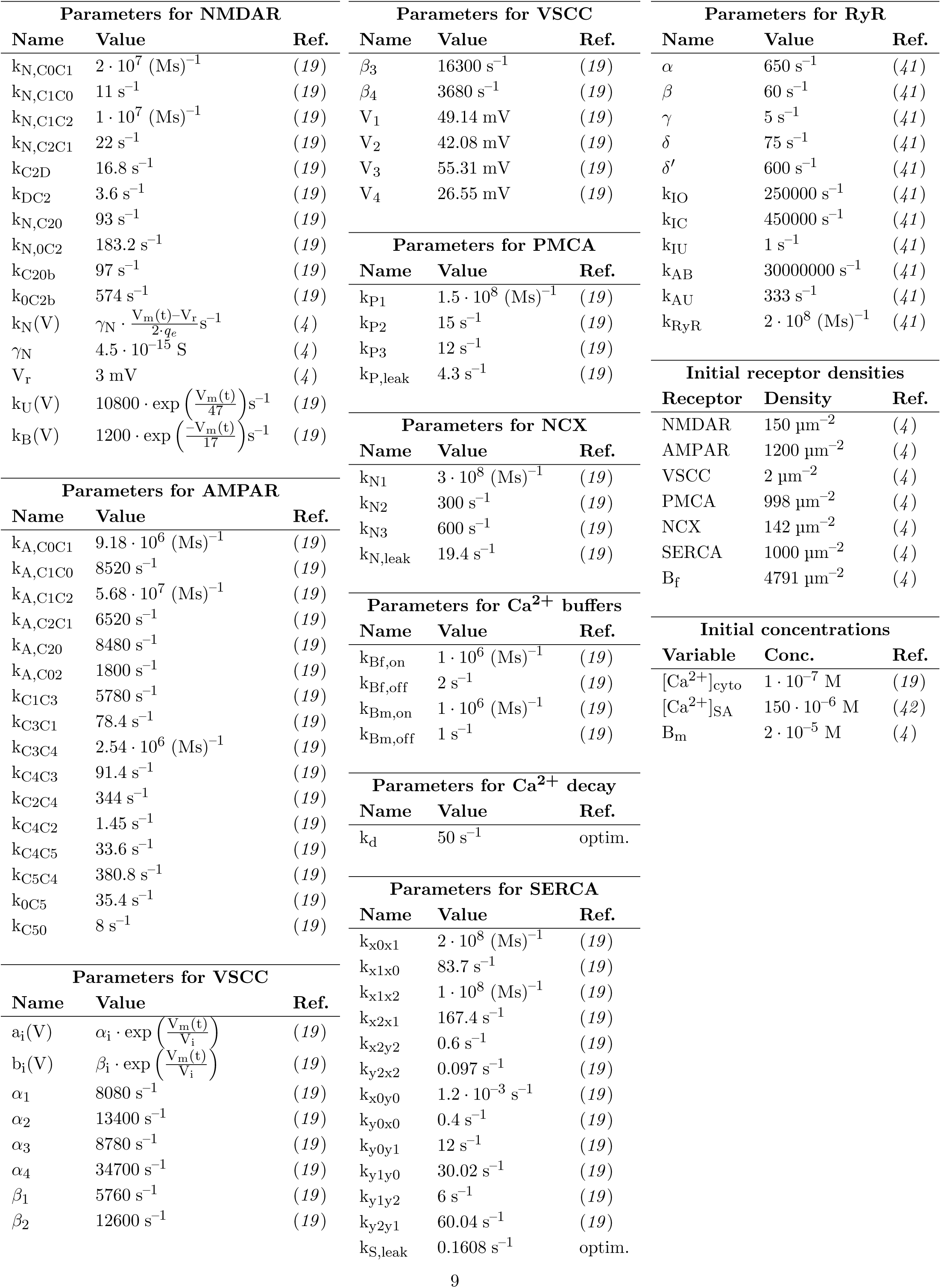
Reaction rates and parameters used for the simulation.

### 4.1 Receptors at the plasma membrane

Most of the plasma membrane transport mechanisms and equations are described in Bell *et al.* (*4*). For completeness, in the following sections, a brief description for each module is given.

#### 4.1.1 NMDAR receptors

As in (*4*) we assume the N-methyl-D-aspartate receptor (NMDAR) model proposed by Vargas-Caballero *et al.* (*35*). The model equations and kinetic rates are obtained from (*35*) and shown in Tables 1 and 2, respectively. A schematic of the NMDAR reaction rate model is given in Figure 4B. The NMDARs are located in the postsynaptic density (PSD) with surface density of 150 µm^−2^. As described in (*4*), Ca^2+^ influx into the cytosol through open NMDARs is determined by the following reaction:

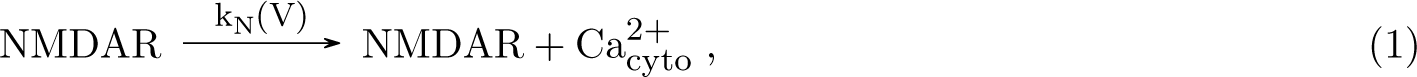

where k_N_(V) is a voltage-dependent rate given by the following expression:

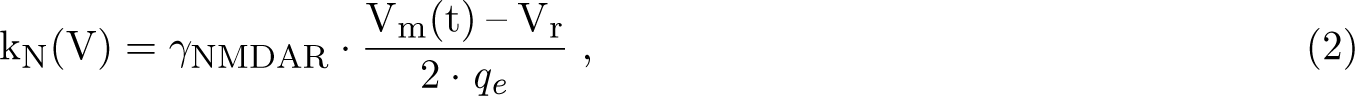

where *q_e_* is the elementary charge. Two further reaction rates, k_U_(V) and k_B_(V), which represent the change between the states NMDAR and NMDAR_B_ are also dependent on the transmembrane potential (Figure 2B) and as in (*19*) are given by the following expressions:

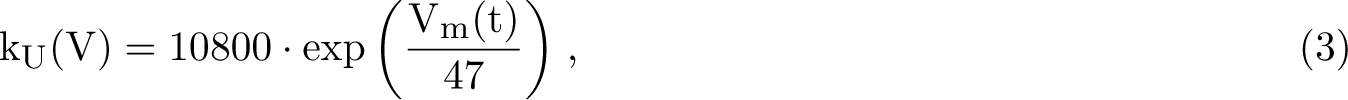

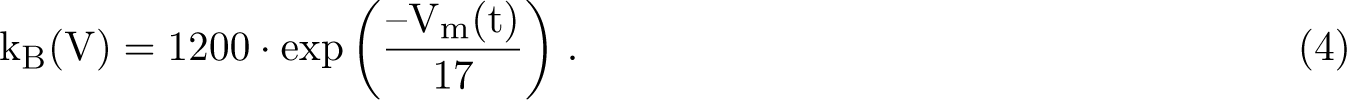

#### 4.1.2 AMPAR receptors

It has been shown that 90% of hippocampal neurons contain GluA2 subunit (*39*), which characterizes the impermeability of *α*-amino-3-hydroxy-5-methyl-4-isoxazolepropionic acid receptors (AMPARs) to Ca^2+^. Therefore, our model does not include Ca^2+^ flux through AMPARs. Rather, AMPARs serve as glutamate buffers to simulate glutamate binding competition with NMDARs. The model of Jonas *et al.* (*36*) was used to simulate the binding of glutamate to AMPAR. We assumed an AMPAR density of 1200 µm^−2^ as described in (*4*, *19*). AMPAR receptors were located only at the PSD.

#### 4.1.3 VSCC receptors

Voltage sensitive Ca^2+^ channel (VSCC) dynamics are based on a 5 state model proposed in (*4*, *19*). The kinetic rate constants within these states are voltage dependent. We use the same density assumption as in (*4*) of 2 µm^−2^ based on measurements in apical dendrites (*40*).

The VSCCs also allow for an influx of Ca^2+^ once they are open which is defined by the following equation:

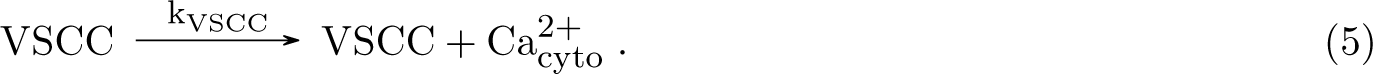

The rate k_VSCC_(V_m_(t)) is membrane potential-dependent and given by the following expression:

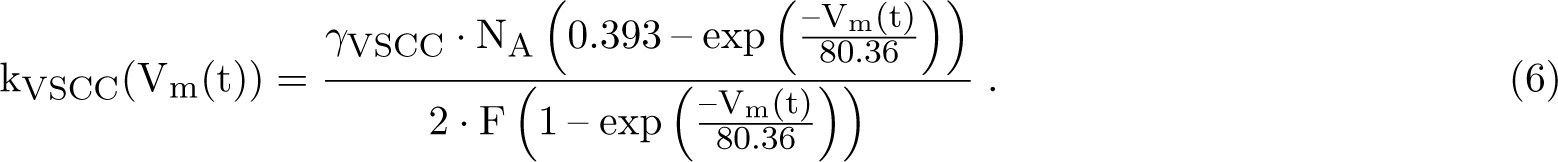

In addition, the reaction rate a_i_(V_m_(t)) and b_i_(V_m_(t)), describing the changes between VSCC states also depend on the transmembrane potential and are given by the following expressions:

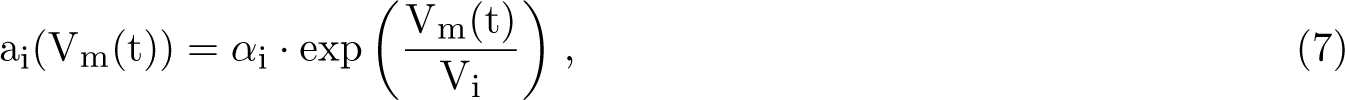

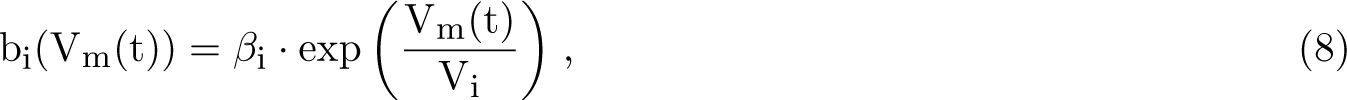

where *α*_i_, *β*_i_ and V_i_ are specific parameters given in Table 2. The VSCCs were assumed to be homogeneously distributed in the spine plasma membrane.

#### 4.1.4 PMCA, NCX and fixed buffers

For modeling PMCA, NCX, and fixed buffers on the plasma membrane, the same equations and parameters as in (*19*) and (*4*) were used. The PMCAs and NCXs were homogeneously distributed along the entire plasma membrane with a receptor density of 998 µm^−2^ and 142 µm^−2^, respectively. The fixed Ca^2+^ buffers were homogeneously distributed in the spine plasma membrane with a density of 4791 µm^−2^.

### 4.2 Equations in the cytosol

We included two reactions occurring in the cytosol. These reactions are Ca^2+^ binding to mobile buffers as described in (*43*) and a decay term for Ca^2+^ to capture other Ca^2+^ buffering events in the cytosol. The decay parameter k_d_ was optimized using the ODE version of the model to match experimental data, see Section 4.4.

### 4.3 Receptors at the smooth ER membrane

In this section we describe the receptor equations for RyRs and SERCA which are the receptors located at the smooth ER membrane. Our model only considers RyRs but not IP_3_Rs based on Spacek *et al.* (*2*), where they showed that RyRs are more common in dendritic spines while IP_3_Rs in the dendritic shaft.

#### 4.3.1 RyR

**RyR dynamics** The presence of ryanodine receptors in neurons is well known (*21*, *44* –*47*). However, to the authors’ knowledge, there is no existing mathematical model that has been specifically designed to match neural RyR experimental data. Therefore, to simulate the effects of RyRs in dendritic spines we used the cardiac model formulated by Tanskanen *et al.* (*41*). This model adaptations allow to use the 4 state model (closed RyR_C_, open RyR_O_, closed inactivated RyR_CI_ and open-inactivated RyR_OI_) from Stern *et al.*(*48*, *49*) in particle-based modelling. It is additionally assumed that each RyR contains 4 Ca^2+^ binding activating sites (RyR(a)) and 4 Ca^2+^ binding inactivating sites (RyR(i)). Thus, each of the original 4 states has different possible substates depending on the number of Ca^2+^ ions bound to the activating and inactivating sites RyR_js_ (a *∼* j_a_, i *∼* j_i_) with j_s_ = *{*C, O, CI, OI*}* representing the 4 states, j_a_ = *{*0, 1, 2, 3, 4*}* representing the number of Ca^2+^ molecules bound to the activating sites, and j_i_ = *{*0, 1, 2, 3, 4*}* representing the number of Ca^2+^ molecules bound to the inactivating sites. Therefore counting states and substates (the substates are given by the number of Ca^2+^ ions bound to the different sites) the modified model contains 100 possible states, instead of 4. The following reactions represent cytosolic Ca^2+^ binding to the activating sites of the RyRs, independently of how many molecules are bound to the inactivating sites. The reaction rate constants are given in Table 2.

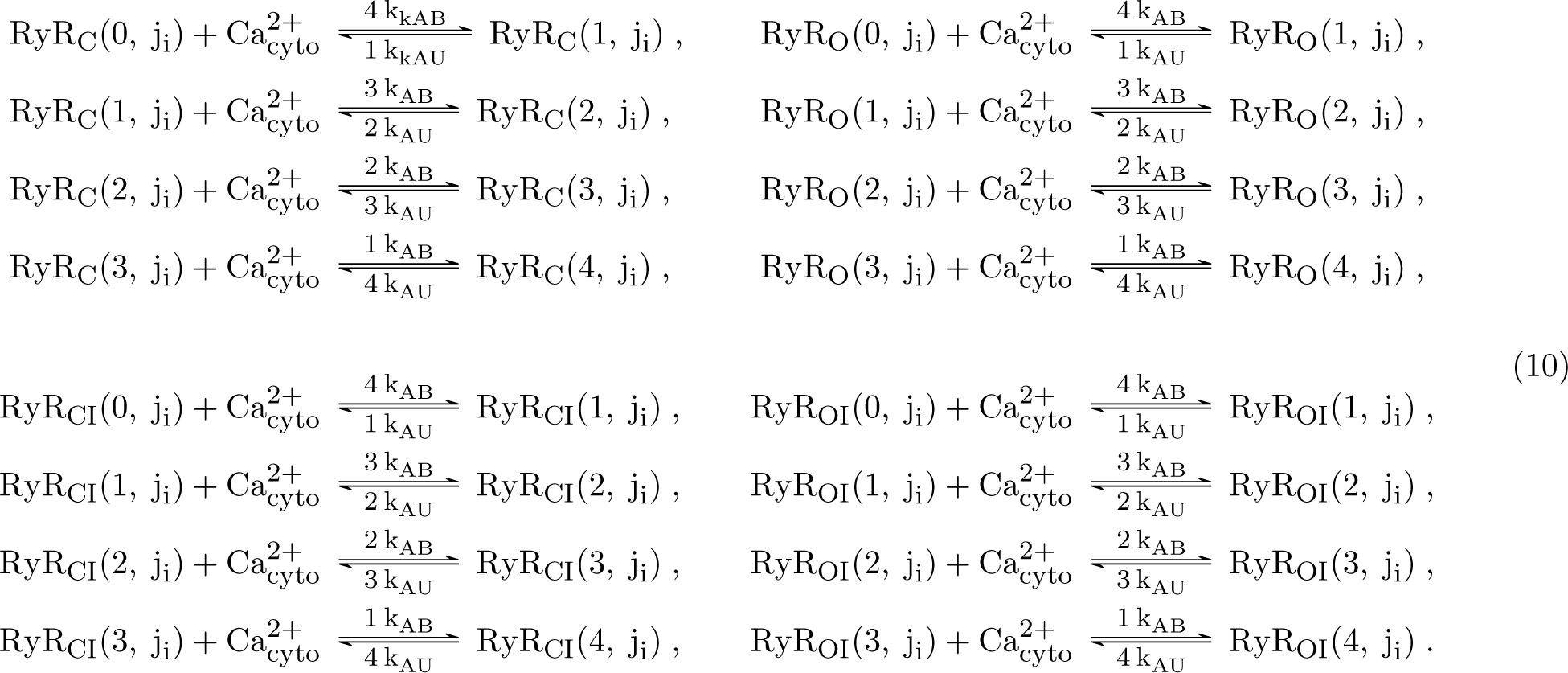

The following reactions represent cytosolic Ca^2+^ binding to the inactivating sites of the RyRs, independently of how many molecules are bound to the activating sites. Note that the binding rates for cytosolic Ca^2+^ binding to the closed states (k_IC_) is different from cytosolic Ca^2+^ binding to the open states (k_IO_).

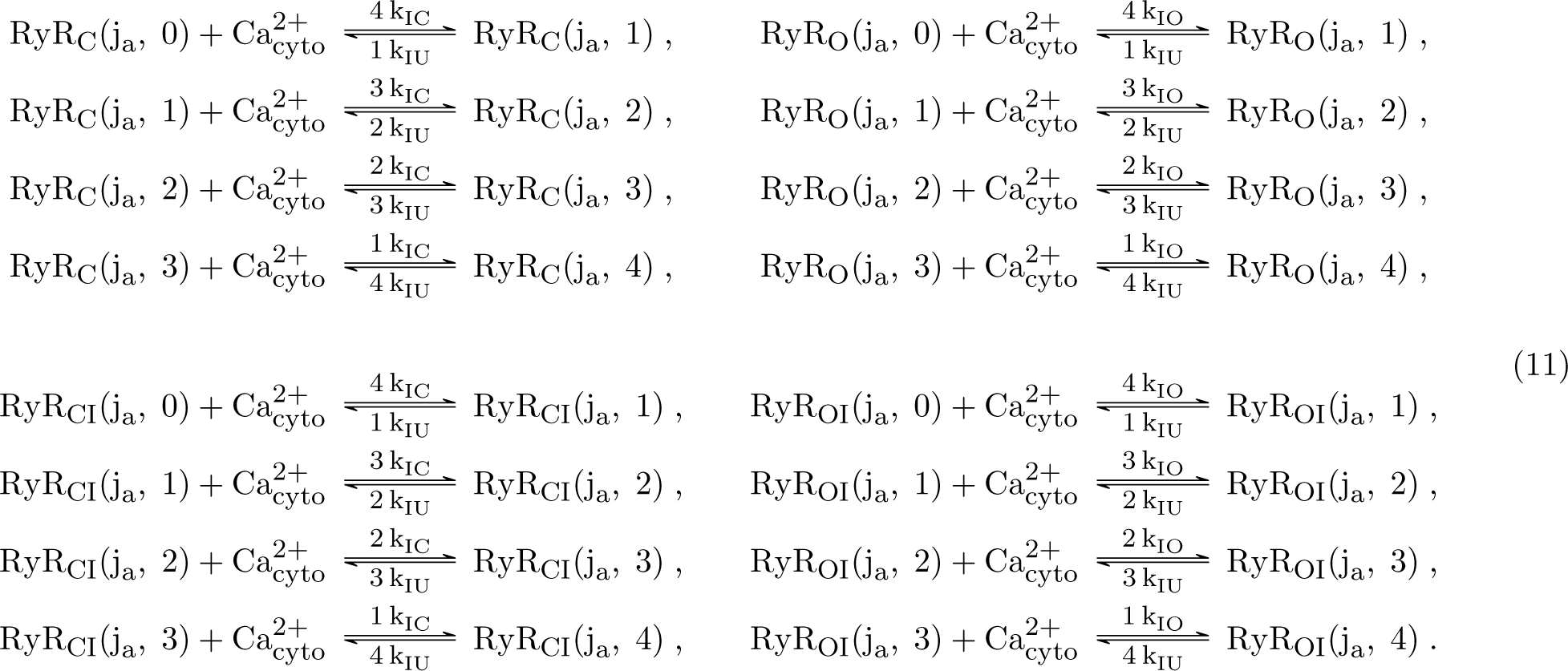

The following reactions represent the transition between closed and open RyRs. The RyRs are always able to close with rate *β*, independently on how many cytosolic Ca^2+^ molecules are bound to the activating or inactivating sites. However for the RyRs to open, at least 4 activation sites must be bound to cytosolic Ca^2+^.

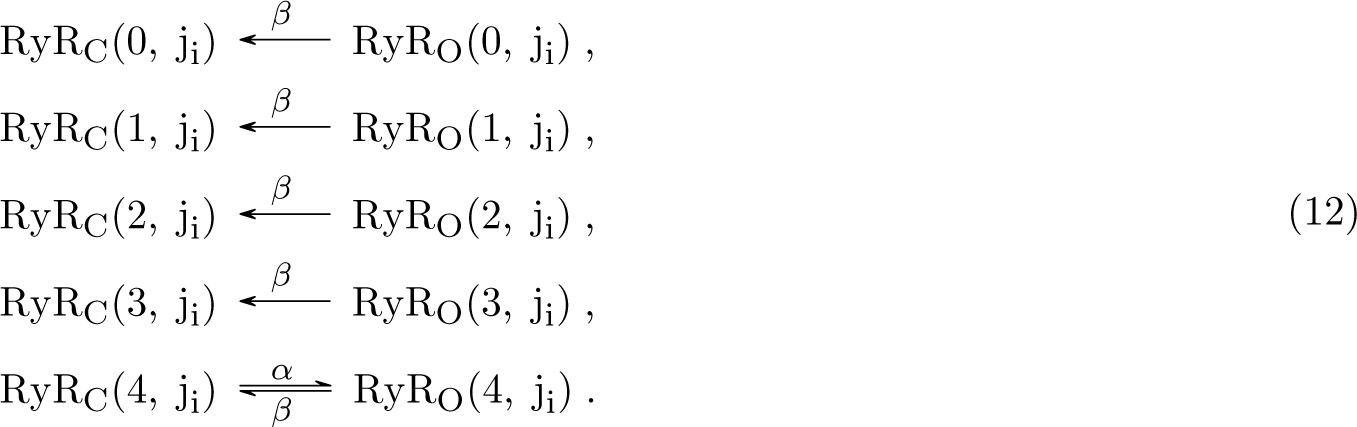

The following reaction governs the transition between closed inactivated and open inactivated RyRs, which occurs independently of the number of cytosolic Ca^2+^ molecules bound to the activating and inactivating sites.

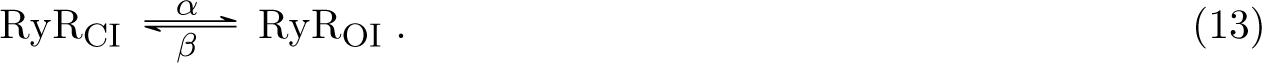

The following reactions represent the transition between closed and closed-inactivated RyRs.

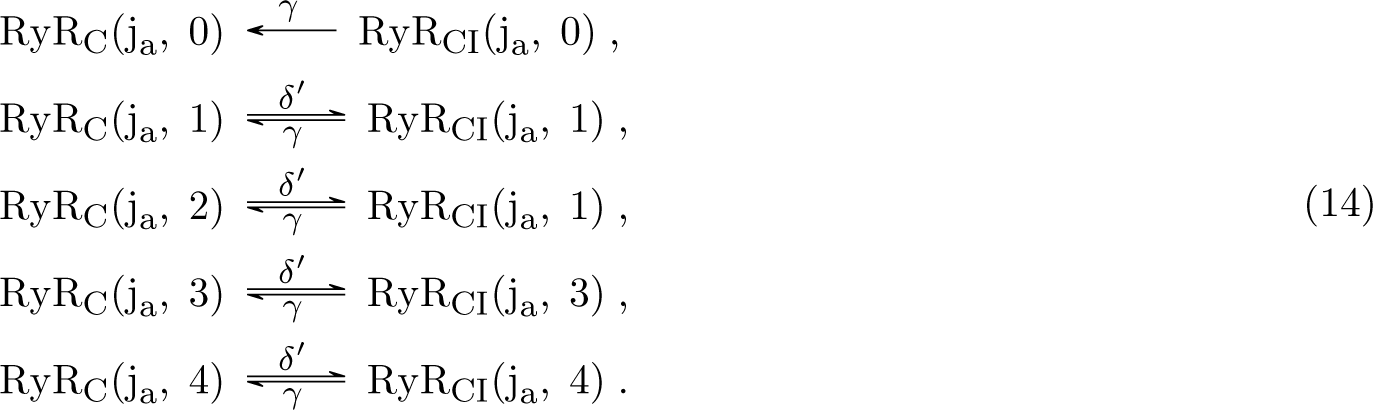

The inactivation rate *δ^′^* is nonzero if at least one Ca^2+^ molecule is bound to an inactivation site. The following reactions represent the transition between open and open-inactivated RyRs.

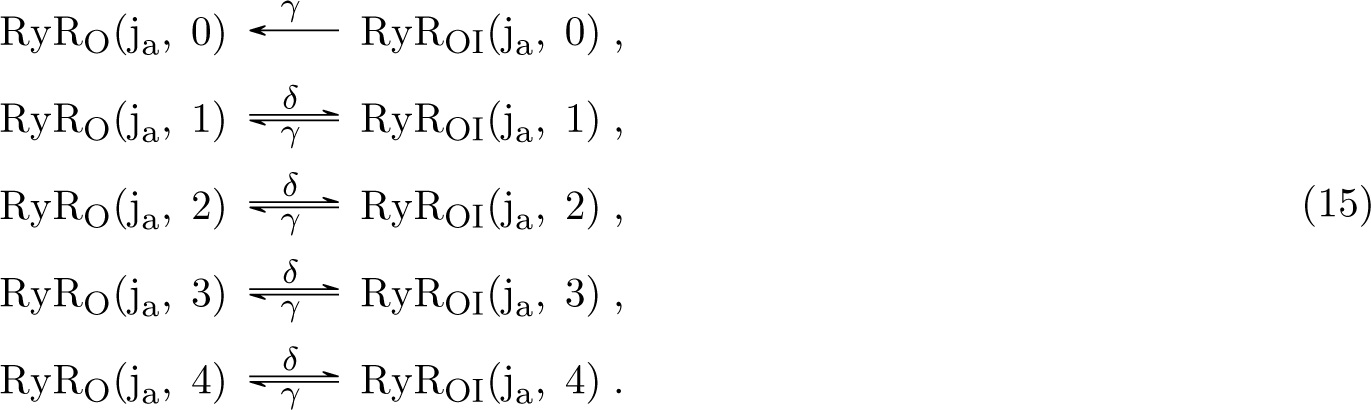

The inactivation rate *δ* is nonzero if at least one cytosolic Ca^2+^ molecule is bound to the inactivation site.

**RyR flux rate** Once the RyRs are opened, Ca^2+^ can efflux from the ER to the cytosol due to the Ca^2+^ concentration gradient. However, we assume a bidirectional reaction since a small influx from the cytosol into the ER is technically also possible. The reaction describing the flux of Ca^2+^ is given by the following expression:

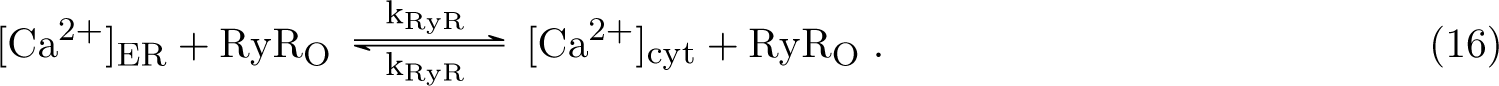

The RyR reaction rate constant k_RyR_ was set to a value of 1.09 *×* 10^9^ m^−1^s^−1^ as suggested by Singh *et al.* (*50*).

**RyR density** We could not find any direct measurements of RyR density in dendritic spines. However, this has been studied in cardyomyocytes using super-resolution imaging (*32*, *33*, *51*). Shen *et al.* measured around 3.4 Ca^2+^ release units per µm^3^ in rat cardiomyocytes, with a Ca^2+^ release unit containing around 20 to 25 RyRs. Furthermore, Hou *et al.* measured a Ca^2+^ release unit density of around 1.1 per µm^2^. To keep our RyR within a similar range to these measured values in the homogeneous cases of both ER geometries we chose to place 60 RyRs and keep this number constant for all distributions.

#### 4.3.2 SERCA

We employ the same SERCA pump model as in (*4*), which was originally formulated in (*19*). The SERCA density at the membrane is 1000 µm^−2^ (*19*). We estimated the SERCA Ca^2+^ leak parameter by optimizing the ODE curve to match the experimental data from (*22*), see Figure S2C.

### 4.4 ODE model

In order to optimize the parameters k_d_ and k_S,leak_, which represent the Ca^2+^ decay and SERCA leak terms respectively, we developed a compartmental ODE version of the model. We used VCell (*52*) to formulate the initial flux equations between compartments and ran the model in MATLAB using the ode solver ode15s. A schematic of the compartments used for the ODE model is given in Figure S2A. The model was first developed without a SA compartment in order to confirm that the ODE model matched the spatial simulations with no ER as described by Bell *et al.* (*4*), see Figure S3A. Next, an inactive ER compartment, including SERCA but no RyRs, was added to the ODE model, which was used to replicate the corresponding MCell simulations, see Figure S3B. Lastly, RyRs were included in the ER compartment of the ODE model. As described in Section 4.3.1 the RyR model we considered contains 100 states. A model with 100 states would be very complex to be implemented in an ODE model. However, this model is a detailed expansion of a four state model originally formulated by Stern *et al.* (*48*, *49*) which allows for particle-based simulations. Therefore, we implemented the four state model within the ODE framework and reliably replicated the dynamics obtained using the spatial stochastic particle-based simulations (Figure S2B). This ODE model was then used to calculate the parameters k_d_ and k_S,leak_ by fitting them to experimental data from Segal *et al.* (*22*) using the MATLAB function for solving non-linear least squares problems lsqnonlin (Figure S2C).

### 4.5 Stimulus and further simulation details

We use the same stimulus approach as in (*4*, *19*), which consists of a plasma membrane depolarization and glutamate release that activates the N-methyl-D-aspartate receptors (NMDARs) (Figure 2). The plasma membrane depolarization includes an excitatory postsynaptic potential (ESPS) and a back propagating action potential (BPAP). The glutamate molecules are released instanteneously at the geometrical center of the PSD.

A further point of discussion is the SERCA distribution. Basnayake *et al.* (*27*) measured SERCA to be mostly located at the synaptopodin sites, which correspond with the sites with highest laminarity. However, by increasing the surface area towards the upper part of the spine apparatus and keeping the SERCA density constant we ensured that a higher number of SERCA was located there.

An initial Ca^2+^ concentration of 150 µM is assumed in the cytosol prior to the stimulus. After glutamate release, Ca^2+^ is released through NMDARs (in the post synaptic density) and VSCCs (in the plasma membrane) due to glutamatergic simulation in the center of the PSD and plasma membrane depolarization. Through the mechanism known as Ca^2+^-induced Ca^2+^ release, further Ca^2+^ ions are then released from the RyRs. At the longitudinal ends of the dendrite, reflective boundary conditions were assumed.

For each simulation setup, we ran 250 realizations in MCell4 (*37*) and calculated mean number of Ca^2+^ ions and standard error of the mean. The Ca^2+^ transient plots are generated using stdshade (*53*) in Matlab version 2022a. In RyR distributions and geometries with less than 50 opened RyRs, we ran 150 or so further simulations. The number of simulations with open RyRs and the total number of simulations is given in Table S1.

### 4.6 Geometries

In order to study the effects of the SA on Ca^2+^ handling we utlized idealized and realistic geometries of both the plasma membrane and the ER. Figure 2C depicts the four geometries used in this work, which are a combination of idealized and realistic plasma membrane and non-laminar SER and laminar SA.

For the idealized plasma membrane geometry, we used the mushroom shaped spine geometry from (*4*, *10*). We expanded this original idealized geometry to include a piece of dendrite in order to calculate how much Ca^2+^ reaches the dendrite in different conditions. The included dendrite consists of a cylindrical structure with a diameter of 1 *µ*m as described in (*54*) and a length of 1 *µ*m (Figure S2D). The ratio between spine apparatus diameter in the dendrite to dendrite diameter was set to 0.166 *µ*m as described in (*55*). A comparison between simulations including a piece of dendrite and the simulations from Bell *et al.* if given in Figure S2E. For the ER geometries used in the geometries with idealized plasma membranes, we also differentiated between non-laminar SER and laminar SA. For the non-laminar case, we used the same approach as in (*4*, *11*) by considering a “spine within a spine”. However, the shape of the mature SA is characterized by its laminar structure consisting of stacked ER cisternae (*18*). Thus, we also designed an idealized SA with laminar stacked discs.

For the realistic spine geometry, we use the plasma membrane and SA segmented and meshed by Lee *et al.* (*56*), originally imaged by Wu *et al.* (*20*) using focused ion beam scanning electron microscopy in the mouse cerebral cortex. The realistic SA from Lee *et al.* (*56*) demonstrates the characteristic laminar shape. The realistic plasma membrane geometry includes two PSDs in contrast to the single PSD of the idealized geometry Figure 4A. For both the idealized and the realistic plasma membrane glutamate was released at the center of the single PSDs at the beginning of the simulation and the values are given in Table 4. In order to systematically study the effect of laminar SA structures, we additionally included an idealized non-laminar ER geometry for the realistic plasma membrane by removing the laminarity and smoothing the ER. The laminar structure of a SA results in a much higher surface area to volume ratio in the ER head. The differences in surface area, volume, and surface area to volume ratio of all spine apparatuses are given in Table 3.

**Table 3:** Surface area, volume, and surface area to volume ratio (SA/V) for the different plasma membranes and ERs used.

|  | **Idealized PM** |  | **Realistic PM** |  |
| --- | --- | --- | --- | --- |
|  | Plasma membrane |  | Plasma membrane |  |
| Surface Area | $7.18 \mu\text{m}^2$ | | $6.63 \mu\text{m}^2$ | |
| Volume | $1.05 \mu\text{m}^3$ | | $0.67 \mu\text{m}^3$ | |
| Ratio SA/V | $6.83/\mu\text{m}$ | | $9.88/\mu\text{m}$ | |
|  | Idealized ER | Laminar SA | Idealized ER | Laminar SA |
| Surface Area | $1.28 \mu\text{m}^2$ | $2.20 \mu\text{m}^2$ | $1.24 \mu\text{m}^2$ | $2.40 \mu\text{m}^2$ |
| Volume | $0.052 \mu\text{m}^3$ | $0.049 \mu\text{m}^3$ | $0.027 \mu\text{m}^3$ | $0.027 \mu\text{m}^3$ |
| Ratio SA/V | $24.47/\mu\text{m}$ | $44.74/\mu\text{m}$ | $45.8/\mu\text{m}$ | $88.47/\mu\text{m}$ |

**Table 4:**
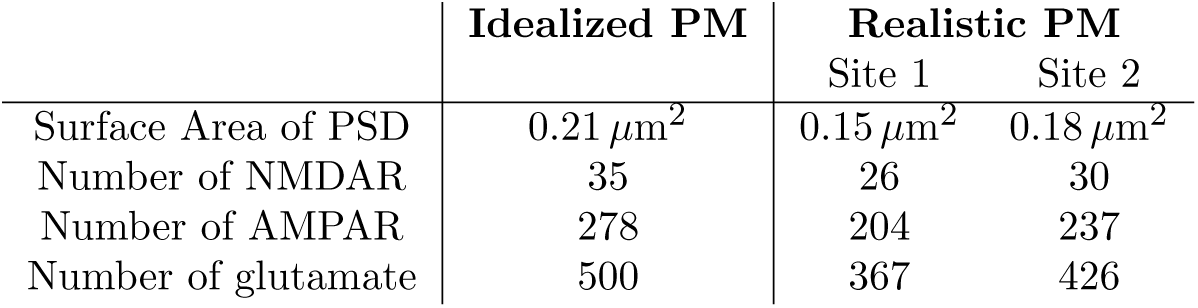
Number of NMDAR, AMPAR and glutamate used for the idealized and realistic plasma membrane simulations. The density of the receptors and the ratio of glutamate to receptors are kept constant in both plasma membrane geometries.

|  | Idealized PM | Realistic PM |  |
| --- | --- | --- | --- |
|  |  | Site 1 | Site 2 |
| Surface Area of PSD | $0.21 \mu\text{m}^2$ | $0.15 \mu\text{m}^2$ | $0.18 \mu\text{m}^2$ |
| Number of NMDAR | 35 | 26 | 30 |
| Number of AMPAR | 278 | 204 | 237 |
| Number of glutamate | 500 | 367 | 426 |

## 5 Results

To understand the relationship between spine geometry, ER geometry, and RyR localization, we used both idealized and realistic geometries from the literature (*10*, *56*) in a spatial, particle-based stochastic model implemented in MCell4 (*37*) and stimulated NMDAR and VSCC activation by including both membrane depolarization and glutamate release. Our simulations reveal the following relationships: RyR opening probability is location-dependent and spine geometry dependent. Importantly, mobile and immobile Ca^2+^ buffers and SERCA can provide some buffering against runaway potentiation of spines even when CICR is activated. For higher Ca^2+^ levels in spine heads, non-laminar SER and high NMDAR opening, with RyR localized at the head are favorable. For higher Ca^2+^ levels in the dendrite, with a potential for spine-to-spine communication, laminar SA and neck localization of RyR is favorable. We elaborate on these findings below.

### 5.1 RyR distribution affects opening probability and Ca^2+^ dynamics in an idealized, non-laminar SER

To study the effect of different spatial distributions of the RyRs within the spine apparatus, we first assumed a smooth, idealized geometry for both the plasma membrane and non-laminar SER (Figure 2C.i). We investigated Ca^2+^ dynamics for four different RyR distributions in this geometry: a homogeneous distribution throughout the whole SER section (Homog. setup), RyRs concentrated at the uppermost part of the spine apparatus close to the PSD (Head setup), RyRs concentrated in the spine neck but completely within the spine volume (Neck setup), and RyRs concentrated in the lowest part of the spine apparatus which overlaps both the spine and dendrite volumes (Low neck setup) (Figure 3A). In all cases, the total number of RyRs was held constant (60 RyRs) (Section 4.3.1) and we conducted *|***N***|* = 250 simulations for each scenario. As we and others have shown before (*32*, *34*), due to the stochastic nature of the model, RyRs open only in a subset of those simulations **n** *∈* **N**. Thus the remaining set **N***\***n** is the set of simulations where RyRs did not open.

**Figure 3:**
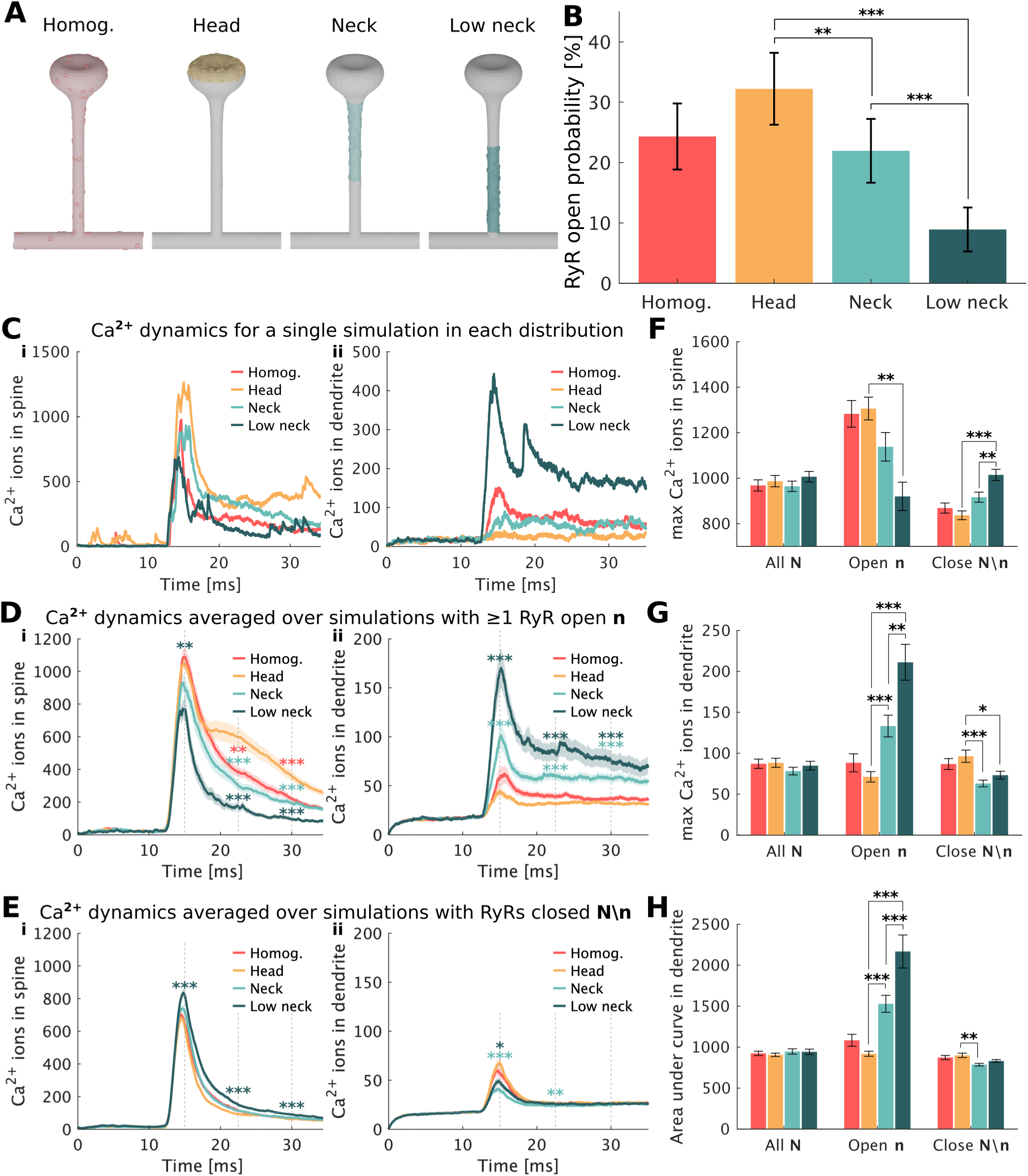
Effect of RyR distribution on Ca^2+^ dynamics in idealized spine and non-laminar SER. **A**: Distributions of RyR considered in idealized SER geometries. **B**: RyR opening probability for each distribution. The bars represent the maximum likelihood estimate of the opening of the RyRs. A total of 250 simulations were run. The p values for the RyR opening probability where calculated from Fisher’s exact test. **C**: Number of Ca^2+^ ions in the spine (left) and dendrite (right) for a single simulation. **D**: Number of Ca^2+^ ions in the spine (left) and dendrite (right) for those simulations containing at least one RyR opened **n**. **E**: Number of Ca^2+^ ions in the spine (left) and dendrite (right) for those simulations containing closed RyRs **N***\***n**. **F**: Max number of Ca^2+^ ions at the spine calculated for each simulation. **G**: Max number of Ca^2+^ ions at the dendrite calculated for each simulation. **H**: Area under the curve for number of Ca^2+^ ions reaching the dendrite (D-E right panels). In plots D-E mean and standard error of the mean are shown. ANOVA and Tukey’s test was performed at times 15 ms, 22.5 ms, and 30 ms with respect to the head distribution. For plots F-H mean and standard error of the mean is shown and the p values are calculated with ANOVA and Tukey’s test. Statistically significant differences are indicated as *p *<* 0.05, **p *<* 0.01 and ***p *<* 0.001.

We observed that the location of the RyRs significantly impacts the probability of RyR opening during a given simulation (Figure 3B). When RyRs are close to the PSD (Head), the opening probability is highest (32.22% *±* 5.95%), and when the RyRs are located further from the PSD, the RyR opening probability decreases (in the lower neck configuration, 8.91% *±* 3.64%). Next, we studied the effect of different RyR distributions along the spine apparatus on the Ca^2+^ dynamics in both the spine and the dendrites (Figure 3C). Figure 3C.i and C.ii show the Ca^2+^ dynamics of a single seed for each RyR distribution in the spine and in the dendrite (Figure S2), respectively. In general, we observe that the spine has higher Ca^2+^ than the dendrite in these simulations.

We next separated the cases where the RyRs open (**n**, Figure 3D) versus closed (**N***\***n**, Figure 3E) to understand how different sources and sinks contribute to Ca^2+^ dynamics. The Ca^2+^ averaged over all simulations **N** is a combination of the subset of simulations with open and closed RyRs weighted by their opening probability and is given in Figure S4. To quantify the differences in the Ca^2+^ ion curves we performed ANOVA and Tukey’s test at timesteps 15 ms, 22.5 ms, and 30 ms between the different distributions and the head distribution. As expected, in the simulations in **n**, the cytosolic Ca^2+^ in the spine is significantly higher than in the case when the RyRs remain closed (compare Figure 3D.i, E.i). When the RyRs are located closer to the PSD (head and homogeneous cases), Ca^2+^ is higher in the spine region and more RyRs tend to open, which through CICR leads to a higher Ca^2+^ peak in the spine (Figure 3D.i). This trend is reversed in the case when the RyRs do not open **N***\***n** (Figure 3E.i) because in this case RyRs merely act as Ca^2+^ buffers. As we will see later, buffering plays an important role in modulating Ca^2+^ dynamics in the spine. Next, we analyzed the differences in Ca^2+^ number in the dendrite based on RyR localization (Figure 3D.ii, E.ii). The amount of Ca^2+^ in the dendrite serves as a marker for synaptic transmission (*57*). In general, less Ca^2+^ ions are observed in the dendrite than in spines due to buffering (compare left and right panels in Figure 3D and E). In the simulations with open RyR **n**, the highest number of Ca^2+^ ions in the dendrite corresponds to the lower neck RyR distribution, followed by the neck distribution. In the case where RyR are localized to the head, a significantly lower amount of Ca^2+^ reaches the dendrite (Figure 3D.ii). As expected, in cases where the RyRs remain closed, the trends for Ca^2+^ in the dendrites are reversed (Figure 3E.ii).

To further analyze these Ca^2+^ trends numerically, we computed the mean and standard error of the mean for the maximum value of Ca^2+^ ions for each single simulation both in the spine (Figure 3F) and in the dendrite (Figure 3G). We observed that for the cases where RyRs were open, high Ca^2+^ in the spine is achieved when RyRs are in the head (1306.3 *±* 50.02 in the head compared to 902.2 *±* 62.52 in the low neck) and high Ca^2+^ reaches the dendrite when the RyRs are located in the neck (71.2 *±* 6.38 in the head compared to 211.2 *±* 21.96 in the low neck). In the closed cases, the opposite trend is observed, with more Ca^2+^ ions reaching the dendrite when the RyR are in the head (96.4 *±* 7.49 in the head compared to 73.3 *±* 4.75 in the low neck) even though there are lower number of ions in the spine (837.0 *±* 19.29 in the head compared to 1014.7 *±* 24.64 in the low neck). The total number of Ca^2+^ ions reaching the dendrite, as indicated by the area under the curve of ions also showed the same trends (compare Figure 3G and Figure 3H). Thus, our results suggest that the location of RyR plays a critical role in determining the spatial impact of RyR opening probability and CICR, thereby affecting spine and dendrite Ca^2+^ concentrations. The differences in dendrite Ca^2+^ concentrations based on RyR location can have implications for neuronal downstream signalling.

### 5.2 RyR localization in the neck increases Ca^2+^ in the dendrite in a realistic plasma membrane geometry

Next, we sought to understand how changing the plasma membrane geometry would affect the opening probability of NMDAR (Figure 4). In the realistic PM geometry, there are two PSDs (Figure 4A) compared to one PSD in the idealized PM geometry. We first simulated one PSD at a time in the realistic geometry (Figure S6). We found that this resulted in few NMDAR opening and therefore decided to stimulate both PSDs simultaneously to ensure sufficient Ca^2+^ influx into the spines. We used a model that allows NMDAR to exist in different states based on Ca^2+^ binding and glutamate binding Figure 4B. Both receptor density and glutamate to NMDAR ratio were kept constant throughout the simulations with realistic and idealized PM as shown in Table 4. We found that the opening probability of NMDAR was significantly higher in the idealized PM (Figure 4C) than in the realistic PM (Figure 4D). We conducted simulations in both geometries with only NMDAR (without AMPAR) to understand why this was happening. When analyzing the different NMDAR states in both the realistic and the idealized plasma membrane, we observed that the activation of C2 state of the NMDAR model was much lower in the realistic plasma membrane configuration compared to the idealized plasma membrane (Figure 4E). Furthermore, when analyzing the average number of NMDAR molecules at state C1 and C2, the idealized PM showed much higher numbers than the realistic PM (Figure 4F and Figure 4G), despite having a lower initial total number of NMDAR receptors at state C0 (Table 4). This indicates that glutamate binding to the NMDARs is reduced in the realistic plasma membrane than in the idealized configuration due to geometrical factors. The PSD areas of the realistic plasma membrane geometry are less concave and regular than the PSD on the idealized plasma membrane. This could justify why in the idealized case the glutamate stays near the PSD area longer, since the concavity “traps” the glutamate molecules. (Figure S7).

**Figure 4:**
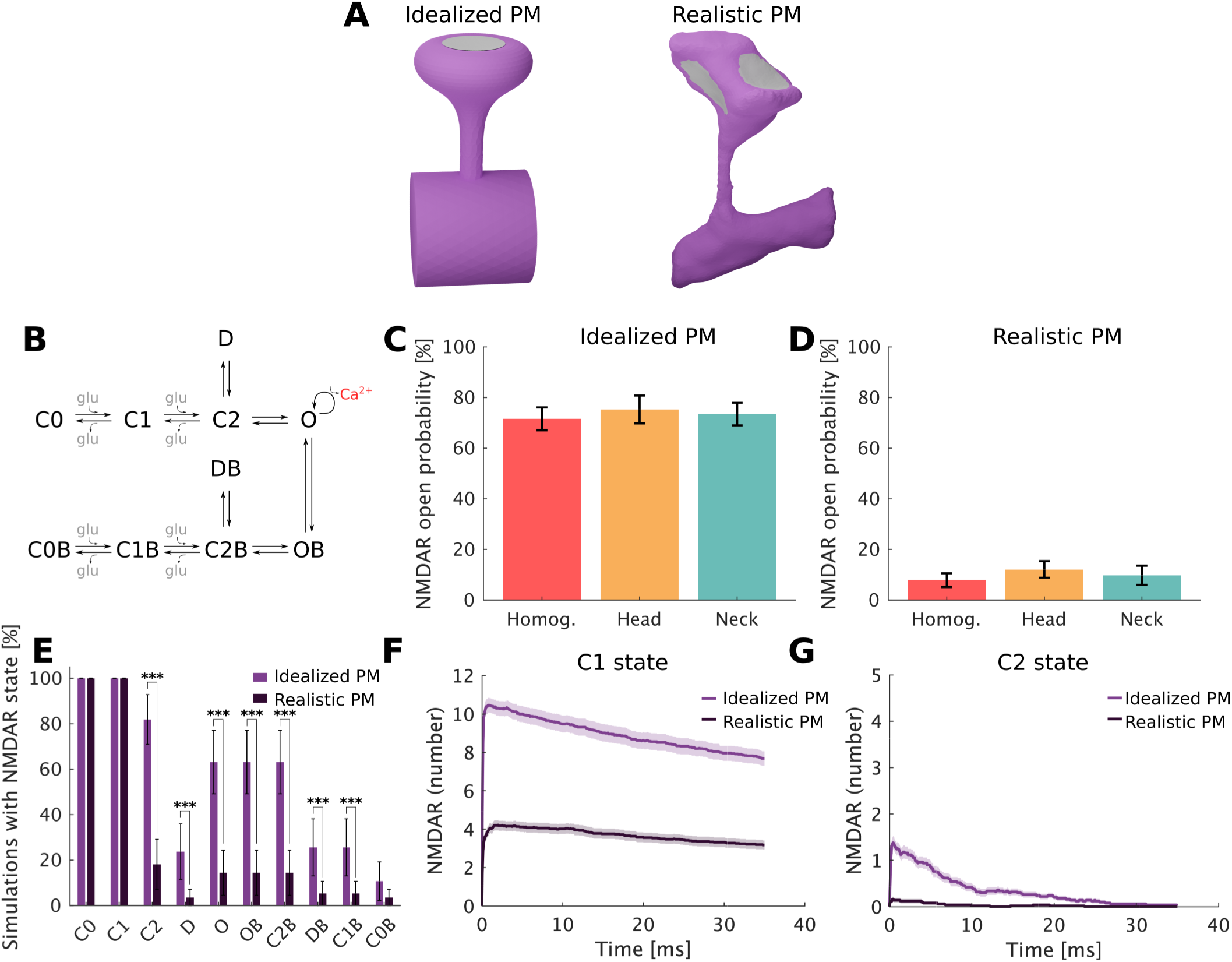
Effect of plasma membrane geometry on NMDAR opening. **A:** Idealized and realistic plasma membrane geometries with the PSD area marked in gray. **B**: Schematic of NMDAR reaction rate model. **C**: NMDAR opening probability for the idealized plasma membrane geometry. **D**: NMDAR opening probability for the realistic plasma membrane geometry. **E**: Probability of each NMDAR state per reaction. The p values were calculated using a two sample t-test. Statistically significant differences are indicated as *p *<* 0.05, **p *<* 0.01 and ***p *<* 0.001. **F**: Averaged number of NMDAR in state C1. **G**: Averaged number of NMDAR in state C2.

Using this realistic PM, we next studied the effects of RyR distribution using a realistic plasma membrane geometry from mouse cerebral cortex spines (*20*, *56*). We smoothed out the measured realistic SA head (Figure 2.iv) to obtain a non-laminar ER head (Figure 2.iii) and kept the realistic ER in neck and dendrites. This allows us to assume a non-laminar SER and run simulations that are comparable to our non-laminar SER and idealized plasma membrane configuration. In this case, we used three RyR distributions: Homog., Head, and Neck (Figure 5A), eliminating “low neck” due to the thin neck geometry. We found that in this hybrid spine apparatus geometry, the opening probability of RyRs *P* (**n**) still strongly depends on the RyR distribution, but is different from the idealized geometry (Figure 5B).

**Figure 5:**
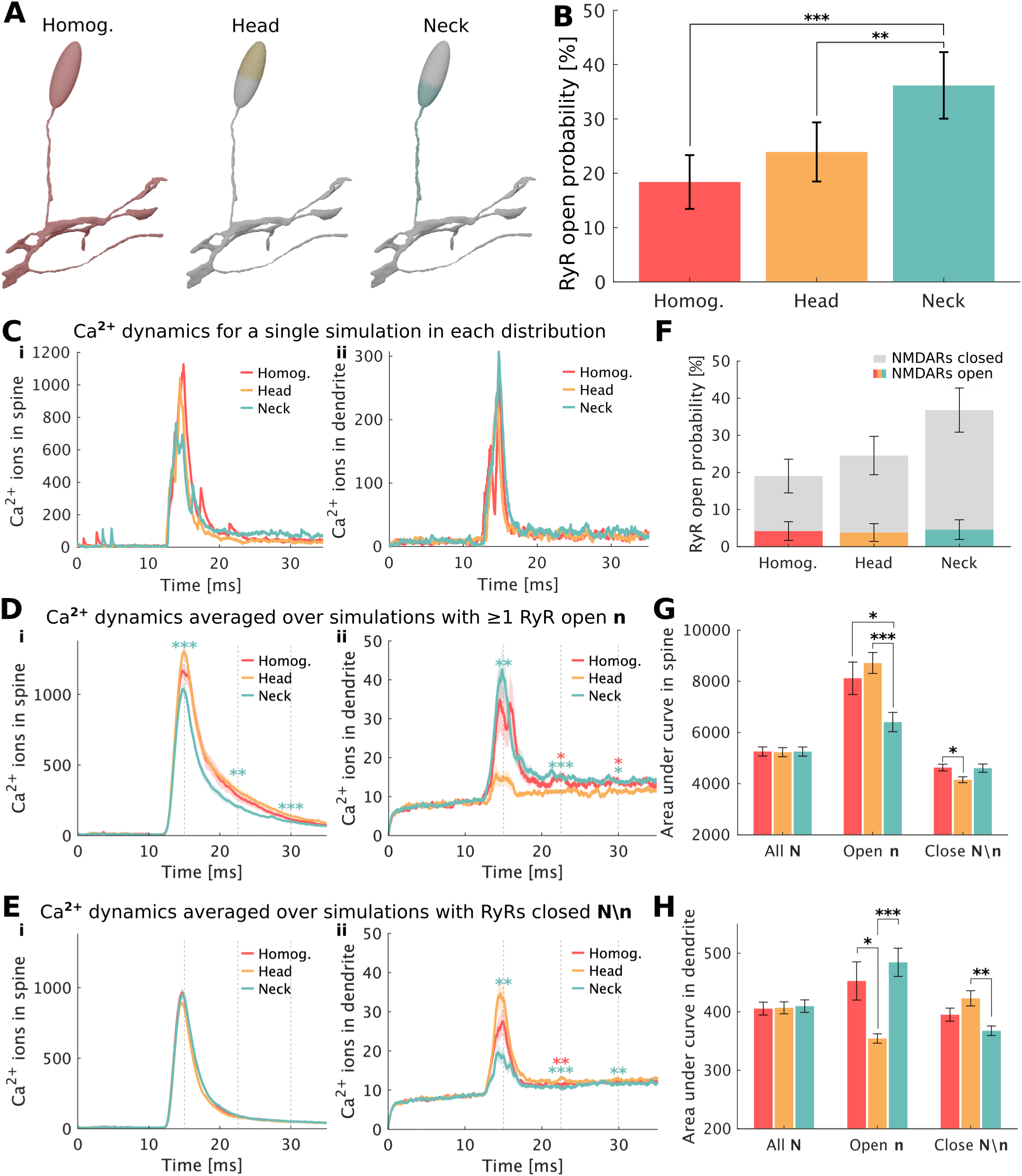
Effect of RyR distribution on Ca^2+^ dynamics in realistic spine and non-linear SER. **A**: RyR distributions considered in the idealized SER geometries. **B**: RyR opening probability for each distribution. The bars represent the maximum likelihood estimate of the opening of the RyRs. The p values for the RyR opening probability where calculated from Fisher’s exact test. **C**: Number of Ca^2+^ ions in the spine (left) and dendrite (right) for a single simulation. **D**: Number of Ca^2+^ ions in the spine (left) and dendrite (right) for those simulations containing at least one RyR opened **n**. **E**: Number of Ca^2+^ ions in the spine (left) and dendrite (right) for those simulations containing closed RyRs **N***\***n**. **F**: RyR opening probability for each distribution including how many of the simulations contain opened NMDARs and how many closed NMDARs. **G**: Area under the curve for number of Ca^2+^ ions in spine. **H**: Area under the curve for number of Ca^2+^ ions reaching the dendrite (D-E right panels). In plots D-E mean and standard error of the mean are shown. ANOVA and Tukey’s test was performed at times 15 ms, 22.5 ms, and 30 ms with respect to the head distribution. For plots G-H mean and standard error of the mean is shown and the p values are calculated with ANOVA and Tukey’s test. Statistically significant differences are indicated as *p *<* 0.05, **p *<* 0.01 and ***p *<* 0.001.

In this case, the highest opening RyR probability 36% *±* 6% was obtained when concentrating the RyRs towards the neck of the spine apparatus, followed by the head (24% *±* 5%) and the homogeneous distribution (18% *±* 5%). Both the homogeneous and head distributions showed a statistically significant lower opening probability when compared to the neck distribution. This is related to the fact that less NMDARs are opened in the realistic PM than in the idealized PM (compare Figure 5F and Figure S8). Therefore, in the realistic case, the Ca^2+^ influx into the dendritic spines is more likely to occur through the VSCCs than through the NMDARs. Since the neck distribution is located closer to the plasma membrane, and the VSCCs are located homogeneously throughout the spine plasma membrane, the neck distribution shows higher RyR opening probabilities.

Figure 5C shows the Ca^2+^ dynamics for a single simulation at the different RyR distributions. In terms of Ca^2+^ dynamics we observe the same trends as in the idealized plasma membrane geometry (Figure 3). For simulations with opened RyRs **n**, high Ca^2+^ in the spine is achieved when RyRs are in the head and high Ca^2+^ reaches the dendrite when the RyRs are located in the neck (Figure 5D, G, H). In the simulations with only closed RyRs **N***\***n**, the opposite trend is observed, with more Ca^2+^ ions reaching the dendrite when the RyR are in the head (Figure 5E.ii, G, H). Higher Ca^2+^ reaches the dendrite in the neck distribution, combined with higher RyR opening probability in the realistic plasma membrane indicate that the RyR neck distribution is the most efficient RyR distribution when accounting for increased spine-to-dendrite communication.

### 5.3 Increased ER laminarity decreases RyR opening probability and Ca^2+^ concentration in the spine

The ER in neurons is highly dynamic and it has been shown to enter and exit spines in an activity-dependent manner (*18*). Recently, Perez-Alvarez *et al.* (*58*) showed that transient ER visits in dendritic spines were mostly synaptopodin negative, whereas 90% of the spines containing stable ER were synaptopodin positive. This suggests that a spine apparatus, whose characteristic laminar SA shape is highly related to synaptodopin (*18*), is more likely to be present in ER stable spines, whereas transient ER visits in spines are more likely to manifest a non-laminar SER. To investigate how the laminarity of the SA would impact Ca^2+^ dynamics, we next simulated ER geometries with laminarities and compared them against non-laminar SER geometries.

We first conducted simulations with the realistic plasma membrane used in Figure 5 combined with the realistic laminar SA measured by Wu *et al.* (*20*). The SA geometry was segmented and meshed by Lee and Laughlin *et al.* (*56*) (Figure 2C.iv). We found that when the ER has a laminar geometry (SA), a higher opening probability is observed when the RyRs are located towards the neck (Figure 6B), similar to observations in the non-laminar SER geometry. We found that the proportion of open RyR simulations containing opened NMDARs was quite low (Figure 6F), as was the case in the realistic plasma membrane with non-laminar SER geometry. Figure 6C shows examples of single-simulation Ca^2+^ dynamics for each RyR configuration both in the spine (Figure 6C.i) and in the dendrite (Figure 6C.ii). In cases where the RyRs open (**n**), the neck setup leads to a significantly lower number of Ca^2+^ ions in the spine (Figure 6D.i) than when the RyRs are localized to the head or when they are homogeneously distributed. However, this effect is reversed when considering how much Ca^2+^ reaches the dendrites, with the head RyR localization having the lowest Ca^2+^ in the dendrite (Figure 6D.ii). This significant difference was also observed when analyzing the total number of Ca^2+^ (area under the curve) in the spine (Figure 6G) and dendrites (Figure 6H). In this case, there are also long time differences in Ca^2+^ in the dendrites in the homogeneous configuration suggesting that there is a cross talk between RyR localization, Ca^2+^ in the spine, and Ca^2+^ in the dendrite. In the simulations where the RyRs stay closed **N***\***n** the differences in Ca^2+^ are small, but more Ca^2+^ reaches the dendrites if the RyRs are placed towards the head of the SA (Figure 6E.ii and Figure 6H). Furthermore, the results for the laminar SA combined with an idealized plasma membrane are given in Figure S5.

**Figure 6:**
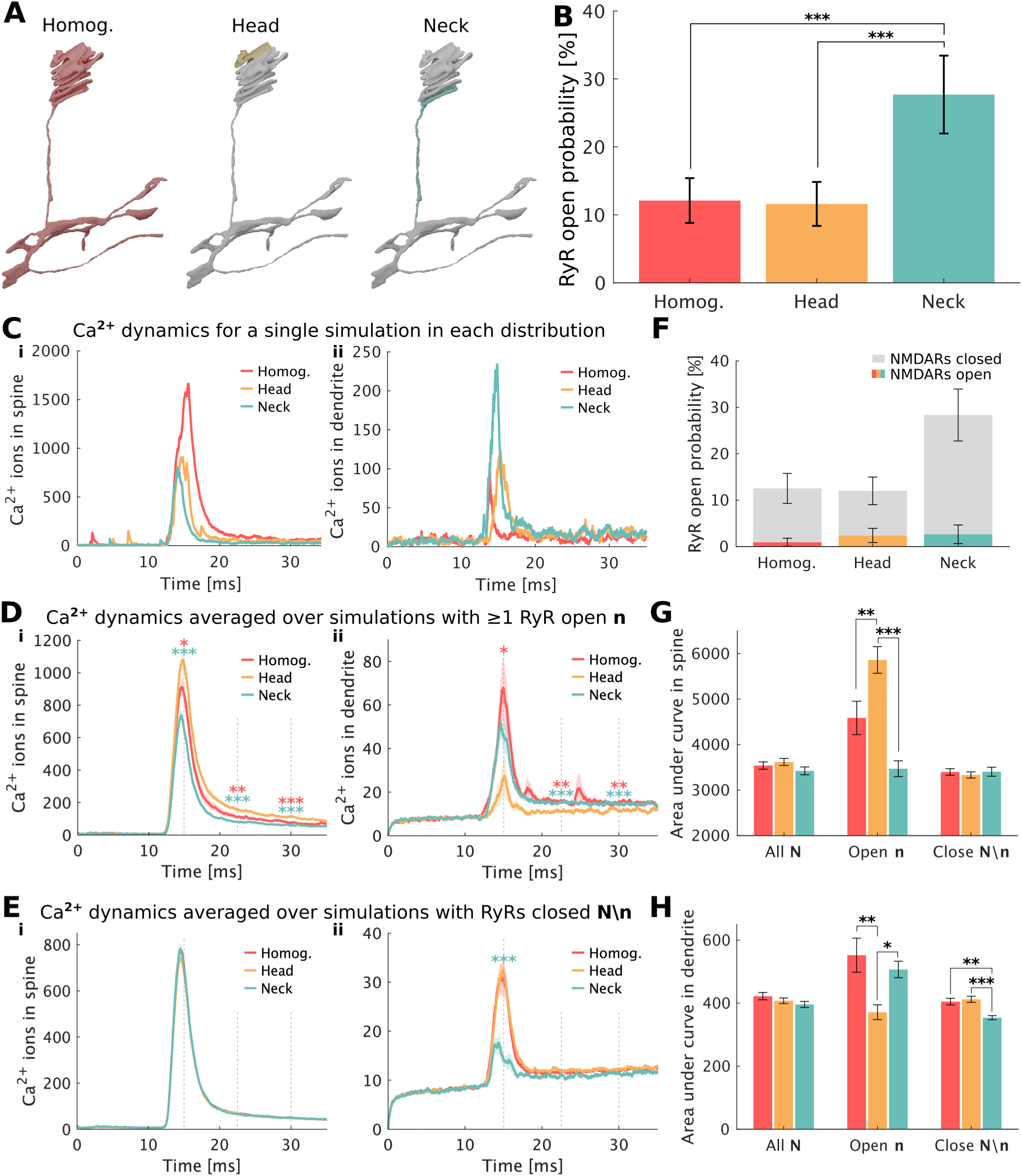
Effect of RyR distribution on realistic plasma membrane and laminar SA. **A:** RyR distributions considered in the realistic SA geometries. **B**: RyR opening probability for each distribution setup. The bars represent the maximum likelihood estimate of the opening of the RyRs. The p values for the RyR opening probability where calculated from Fisher’s exact test. **C**: Number of Ca^2+^ ions in the spine (left) and dendrite (right) for for a single simulation. **D**: Number of Ca^2+^ ions in the spine (left) and dendrite (right) for those simulations containing at least one RyR opened **n**. **E**: Number of Ca^2+^ ions in the spine (left) and dendrite (right) for those simulations containing no open RyRs **N***\***n**. **F**: RyR opening probability for each distribution including how many of the simulations contain opened NMDARs and how many closed NMDARs. **G**: Area under the curve for number of Ca^2+^ ions in spine. **H**: Area under the curve for number of Ca^2+^ ions reaching the dendrite (Fig. D-E right panels). In plots D-E mean and standard error of the mean are shown. ANOVA and Tukey’s test was performed at times 15 ms, 22.5 ms, and 30 ms with respect to the head distribution. For plots G-H mean and standard error of the mean is shown and the p values are calculated with ANOVA and Tukey’s test. Statistical significant differences are indicated as *p *<* 0.05, **p *<* 0.01 and ***p *<* 0.001.

Non-laminar tubules of SER enter dendritic spines (*18*), while in more mature and stable spines the characteristic laminar SA structure is observed (*2*, *58*). This physiological differences between nascent ER and mature SA motivated us to study whether there are any differences in spine-to-dendrite communication based on geometrical properties of the ER in the dendritic spines. To fully understand the effect of laminarity, we directly compared simulations of laminar and non-laminar ER structures (Figure 7) for different PM geometries. In all cases, the RyR number was kept constant, as was the SERCA density. We found that the RyR opening probability was lower in the laminar case than in the non-laminar case for all distributions of RyR (Figure 7A). As a result, the maximum Ca^2+^ in the spines was also lower in the laminar case when compared to the non-laminar case (Figure 7B.i, ii and Figure 7C). We further investigated the different sources and sinks of Ca^2+^ in the spine. One of the main changes that comes with adjusting SA laminarity is surface area of the membrane. When the surface area of the SA changes, the number of SERCA pumps changes because we maintained the density constant based on the fact that SERCA was observed on experimental observations to be placed towards the head, where the laminarity of the SA is located (*17*, *27*). As a result, more Ca^2+^ is pumped back into the SA in the laminar case, whereas increased Ca^2+^ in the spine is observed in the non-laminar case. To understand these effects, we ran simulations on non-laminar and laminar ER containing only SERCA and no RyR and the same effects were observed (Figure S9A). However, there is no change in the Ca^2+^ in the dendrite because of Ca^2+^ buffering by mobile and immobile buffers (Figure S9B). Furthermore, when the number of SERCA, rather than the density is kept the same in the laminar and non-laminar geometries, we found that these spatial effects were eliminated (Figure S9C). Thus, our analysis suggests that the complex Ca^2+^ dynamics are not simply modulated by geometric factors such as surface-to-volume ratio but rather the balance between sources and sinks on different surfaces and volumes. When we now compare the Ca^2+^ in spines with either idealized PM or realistic PM, we find that spine Ca^2+^ is higher in the non-laminar case than in the laminar case (Figure 7C). Conversely, when RyRs are concentrated at the neck, the laminar SA structure leads to non statistically significant more Ca^2+^ reaching the dendrite compared to non-laminar geometries (Figure 7D). Thus, our simulations predict that laminar SA structure is effective at increasing Ca^2+^ concentration in the dendrite. This may have potential consequences for how spines communicate with one another.

**Figure 7:**
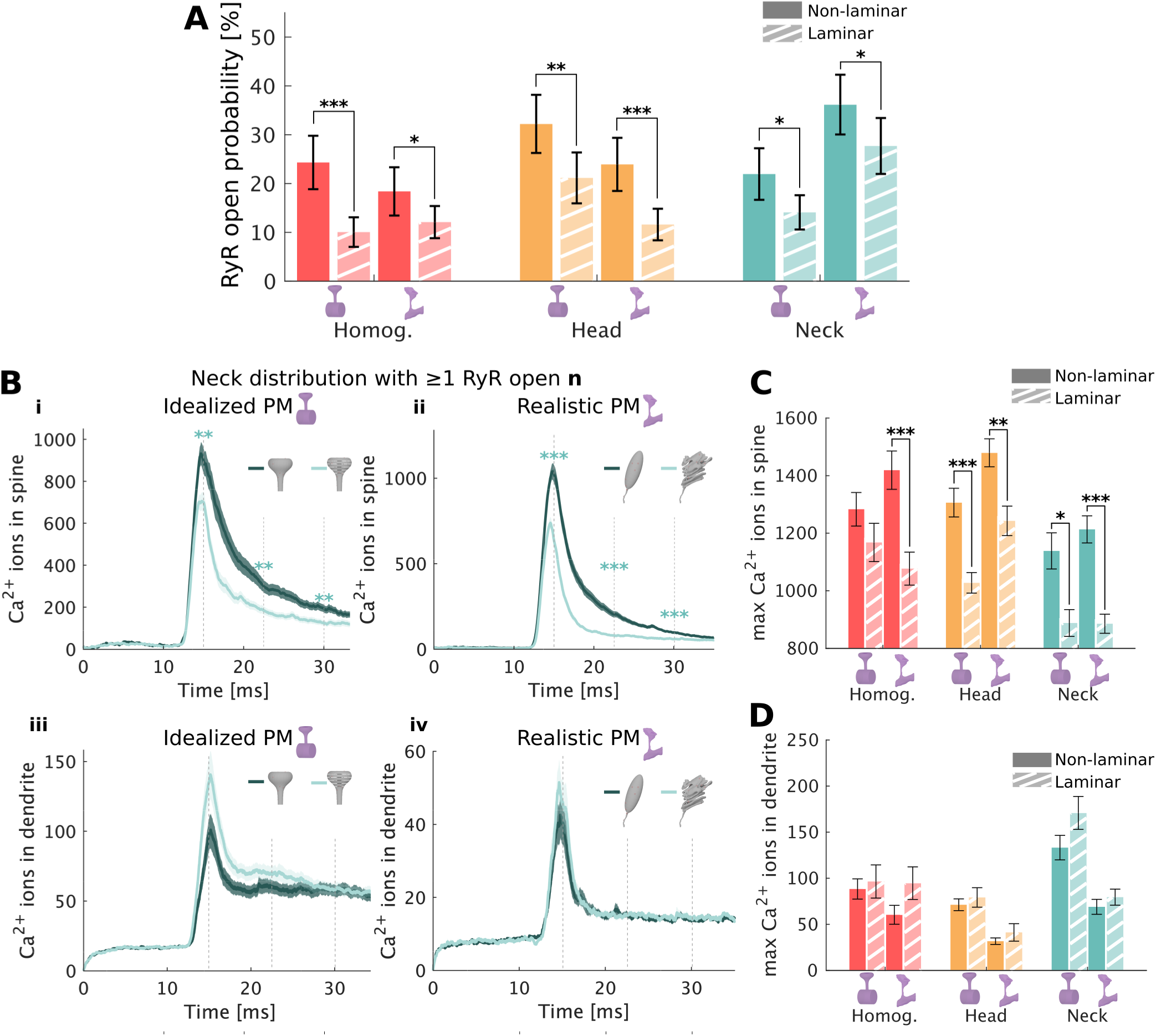
Comparison between a non-laminar SER and a laminar spine apparatus for both plasma membrane geometries. **A:** RyR opening probability comparing non-laminar SER and laminar SA for both plasma membrane geometries. **B**: Comparison between number of Ca^2+^ ions assuming non-laminar and laminar ER in the spine (i and ii) an in the dendrite (iii an iv) for both idealized (i an iii) and realistic plasma membrane geometries (ii and iv). Mean and standard error of the mean is shown. A two sample t-test was performed at times 15 ms, 22.5 ms, and 30 ms. **C**: Maximum number of Ca^2+^ ions at the spine calculated for each simulation. **D**: Maximum number of Ca^2+^ ions at the dendrite calculated for each simulation. For plots C-D mean and standard error of the mean is shown and the p values are calculated with a two sample t-test. Statistical significant differences are indicated as *p *<* 0.05, **p *<* 0.01 and ***p *<* 0.001.

## 6 Discussion

In this work, we sought to decipher the role of the ER in regulating spine and dendrite Ca^2+^ dynamics. However, this is a complex problem where one needs to consider the geometry of the spine and the geometry of the ER itself. In some nascent spines, transient SER are dominant while more mature spines tend to contain a specialized form of the SER, the SA. The precise functional differences between the SER and the SA remain elusive. Morphological changes in the SA have been observed after the induction of long-term potentiation (*16*, *59*), and ER disturbances have been seen in a number of neurodegenerative disorders (*60*).

Here, we implemented a stochastic particle-based model of Ca^2+^ transients in dendritic spine to understand how RyR distributions on the SER and SA, as well as ER laminarity, affect Ca^2+^ dynamics. The model was implemented on an idealized spine plasma membrane as well as on a realistic spine reconstructed from hippocampal rat neurons (*20*, *56*). First, to disentangle the contribution of membrane geometry on Ca^2+^ dynamics we use both an idealized spine plasma membrane and an idealized SER. We found that RyR distribution impacts the likelihood of RyR opening and Ca^2+^ dynamics. In particular, our model predicts that distributing the RyRs towards the neck of the ER, as they have been suggested to be preferentially located (*27*), results in a higher peak Ca^2+^ and total number of Ca^2+^ ions reaching the dendrite if the RyRs open, whereas concentrating RyRs in the head results in higher spine Ca^2+^ but a lower number of ions reaching the dendrite. Our results on the efficiency of localizing the RyRs towards the neck match previously reported results in a computational study replicating a Ca^2+^ uncaging stimulus (*27*). A larger amount of Ca^2+^ reaching the dendritic shaft could have downstream consequences activating other organelles in the dendrite as well as in spine-to-spine communication or spine clustering (*61*). This becomes especially important when we consider the fact that SA is found in more mature spines which tend to be clustered close to smaller spines. We found that the trends in Ca^2+^ are consistent and statistically significant throughout the different geometries distributing the RyRs towards the neck or head.

Interestingly, opposite trends were observed on simulations where the RyRs stayed closed, with more Ca^2+^ reaching the dendrite if the RyRs were located towards the head of the ER. These differences between open and closed RyRs are related to the fact that RyRs are activated through Ca^2+^-induced Ca^2+^ release. Thus, RyRs act as a Ca^2+^ source when activated, but as a Ca^2+^ buffer if they remain closed. A further takeaway is the effect of the plasma membrane geometry on RyR opening probability. The shape of the plasma membrane has an effect on glutamate binding to the NMDARs. While the simulations with opened RyRs in the idealized plasma membrane contained mostly opened NMDARs, this was not the case in simulations in the realistic plasma membrane, where most of the RyR activation occurred *via* Ca^2+^ influx through the VSCCs. As a result the realistic plasma membrane showed the highest RyR opening probability when placing the RyRs towards the neck of the ER. Imaging experiments have shown that VSCC-mediated Ca^2+^ influx dominates the Ca^2+^ dynamics in dendritic spines (*62* –*64*), as was the case in simulations with realistic plasma membrane geometries. The fact that a higher RyR neck opening probability is achieved when VSCC Ca^2+^ influx dominates further emphasizes the importance of RyR localization towards the neck to improve spine-to-dendrite communication.

Furthermore, the laminarity of the SA affects both the opening probability of the RyRs and the Ca^2+^ transients. In both idealized and realistic plasma membranes we observed that ER laminarity (a common characteristic of mature SA geometries) leads to a lower RyR opening probability and a slightly higher, statistically non-significant, number of Ca^2+^ ions reaching the dendrite. This result is an outcome of the complex interplay between geometry, SERCA, and Ca^2+^ buffers. Although a detailed understanding of the mechanisms of the ER anchoring to the dendritic spine and the formation of SA still remains elusive (*18*, *65*), recent studies have shown that spines with transient ERs were mostly synaptopodin negative, while synaptopodin was detected in 90% of stable ER spines (*58*). This, together with the fact that synaptopodin contributes to the characteristic laminar disc structure of SA (*65*), suggests that laminar SA are more related to stable ER spines than non-laminar ER geometries. The outcomes of our study give further insight into the effects and differences between laminar and non laminar ER structures on Ca^2+^ transients, and provide evidence that mature, laminar SA structures are more likely to facilitate efficient spine-to-dendrite communication.

One limitation of our study is related to the gating rates of the RyR model used. The RyR model was taken from Tanskanen *et al.* (*41*), which was designed to replicate RyR dynamics in cardiac myocytes. However, due to the difficulty of measurements in small volumes of single spines (*66*) we could not find any experiments related specifically to the gating of RyRs in these structures. While we acknowledge that our simulations might not match the quantitative RyR opening probabilities measured in spines, our results give insights into the qualitative trends that different RyR distributions have on RyR opening probabilities. To account for this unknown related to realistic probability of RyR opening during a given stimulus event, we have divided the simulation result set **N** into two subsets **n** and **N***\***n**, those containing at least one open RyR throughout the simulation and that only have closed RyRs, respectively. This allows us to understand the effect of RyR opening independent of the RyR opening probability. Previous experimental studies have reported a contribution of RyR mediated Ca^2+^ release to long-term potentiation (LTP) (*67* –*69*). Furthermore, a recent study has proposed that RyRs amplify Ca^2+^ signals generated at the dendritic spines following stimulation of postsynaptic NMDA receptors (*70*, *71*). Therefore, it is likely that the quantitative opening probability of RyRs in spines is higher than those measured in our simulations. Nevertheless, our results give new insights in understanding the effect of an active ER and RyRs on spine-to-dendrite Ca^2+^ communication.

We were unable to find any experimental data regarding the direct measurement of Ca^2+^ concentration within the SA. We chose a ER Ca^2+^ concentration of 150 µM as used in a previous computational model on cerebellar Purkinje cells (*42*) which used a value measured in mouse pancreatic acinar cells (*72*). In cardiomyocytes, however, sarcoplasmic reticulum concentrations have been measured up to 1.2-1.8 mM (*73*). Due to the high variability in ER Ca^2+^ concentrations, further experimental investigation is required to constrain the values for Ca^2+^ concentration in dendritic spines. Furthermore, as shown by Dittmer *et al.* (*74*) increased ER content could increase the Ca^2+^ storage capacity of spines, which is associated to large storing capacity of large spine volumes as observed during LTP.

Computational models help explore biophysical processes in dendritic spines over several timescales going from a few milliseconds (*4*, *27*, *29*) to several minutes (*12*, *75*, *76*) of Ca^2+^ signaling. Our model shows how the interplay, between NMDAR, VSCCs, ER geometry, RyR location, Ca^2+^ buffers and SERCA modulate Ca^2+^ dynamics and can have effects on downstream signaling and spine-to-dendrite communication. We note that our results are focused on early stage membrane depolarization stimulus and the impact of these effects should be studied for longer events, and high frequency stimulation of the spines (*77*, *78*). It has been shown that Ca^2+^ influx through the L-type Ca^2+^ channels leads to further release of ER Ca^2+^ (*79*) at the timescale of several seconds. Furthermore, it has been recently reported that Ca^2+^ efflux from CICR mechanisms activates stromal interaction molecule 1 (STIM1) feedback inhibition of L-type Ca^2+^ channels (*74*, *79*). Therefore, future studies with longer simulation events should include the crosstalk between ER, L-type Ca^2+^ channels and STIM1 to understand how RyR distribution and ER laminarity might affect this mechanism. Future studies should also investigate how our results affect other neuronal downstream signaling, such as the influence of Ca^2+^ on gene transcription at the nucleus (*14*) and other signaling pathways.

## Acknowledgements

We thank Emmet Francis and Ali Khalilimeybodi for their proofreading and their valuable comments. VCell is supported by NIH Grant Number R24 GM134211 from the National Institute for General Medical Sciences. MCell development is supported by the NIGMS-funded (P41GM103712) National Center for Multiscale Modeling of Biological Systems (MMBioS). The simulations were in part performed on resources provided by Sigma2 - the National Infrastructure for High Performance Computing and Data Storage in Norway. MHM and KJM are supported by the Simula-UCSD-University of Oslo Research and PhD training (SUURPh) program, an international collaboration in computational biology and medicine funded by the Norwegian Ministry of Education and Research. FH is supported by NIH grant NINDS NS115947. This work was supported in part by Air Force Office of Scientific Research (AFOSR) MURI FA9550-18-1-0051 to PR.

## 7 Supplemental material

**Figure S1:**
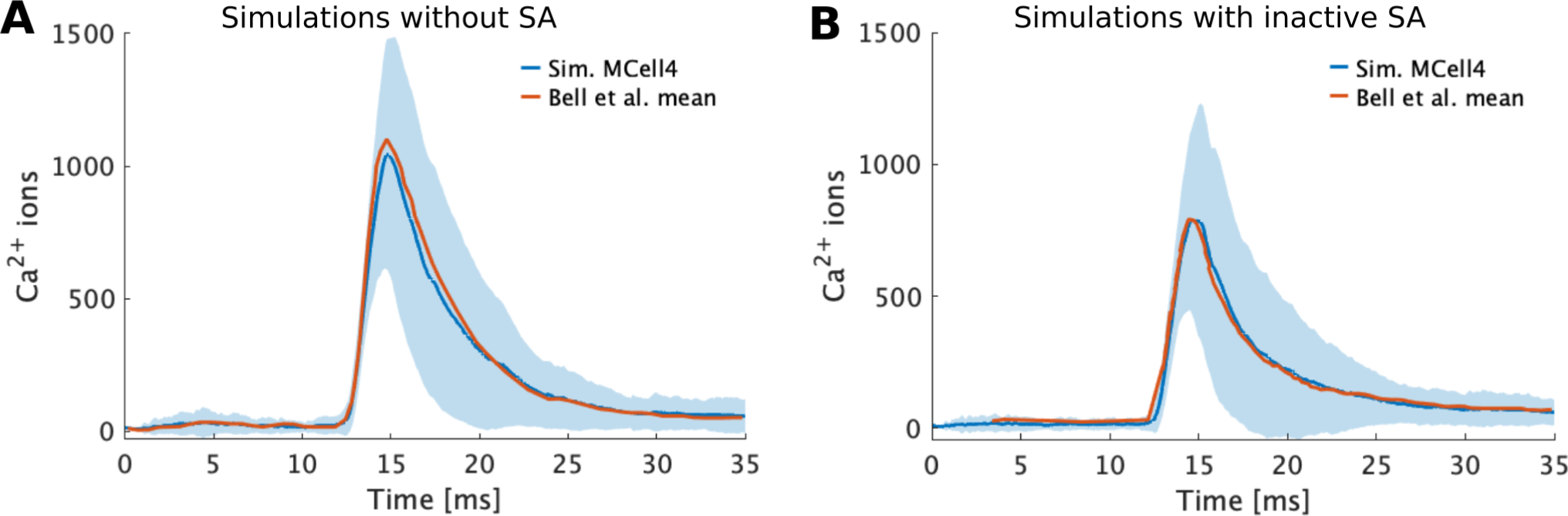
Comparison of model implementation in MCell3 and MCell4. **A**: Simulations were ran without spine apparatus in MCell4 and are compared to the mean value from Bell *et al.* (*4*), where they used MCell3. **B**: Simulations were ran with an inactive spine apparatus (only including SERCA) in MCell4 and are compared to the mean value from Bell *et al.* (*4*), where they used MCell3. In both figures mean and standard deviation of the number of Ca^2+^ ions is shown.

**Table S1:**
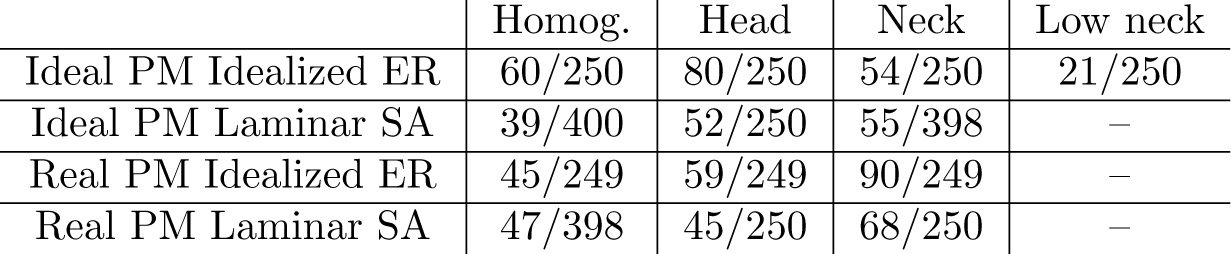
Number of opened RyRs and number of total simulations ran.

**Figure S2:**
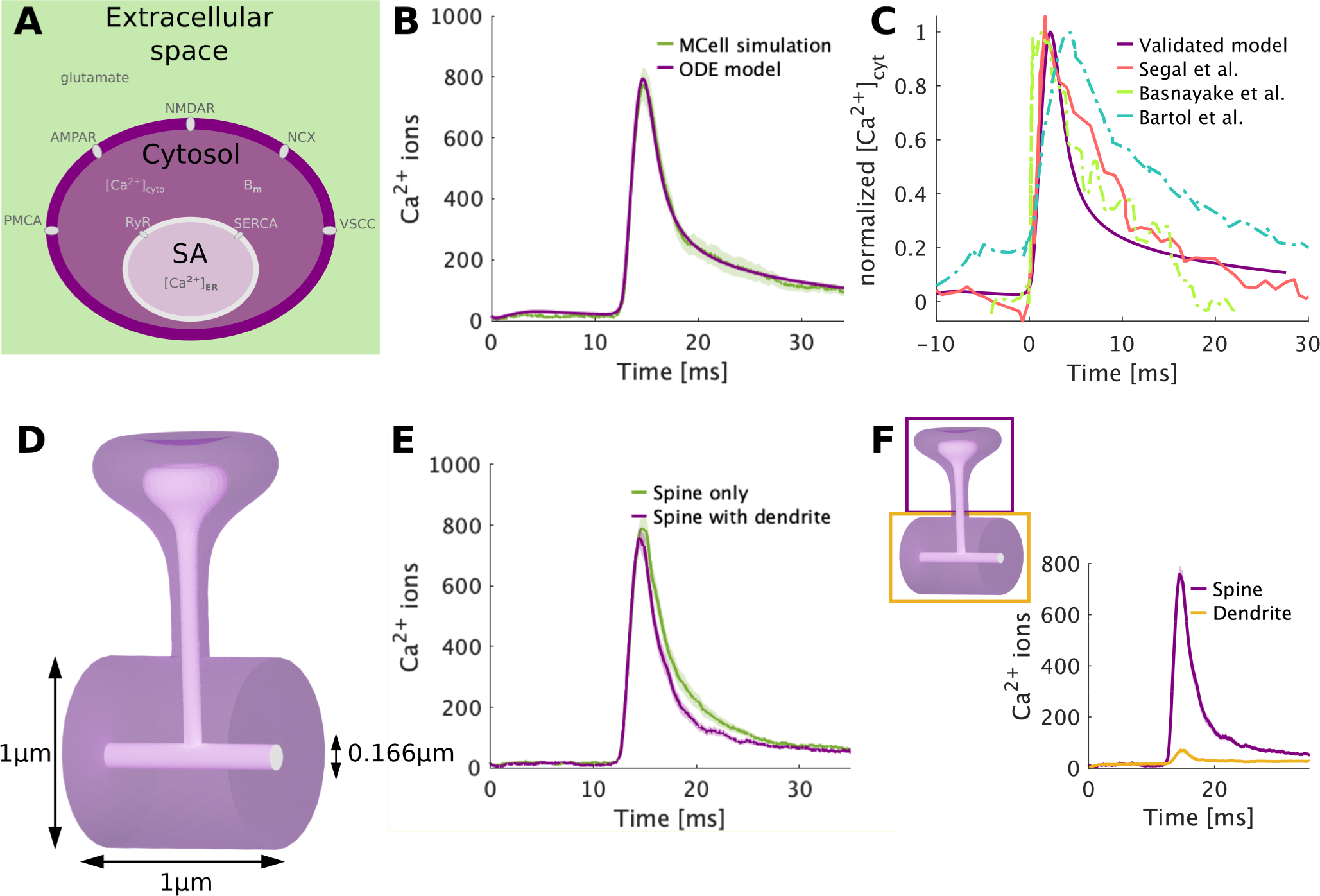
Model calibration. **A:** Schematic of the compartmental ODE model used for the parameter estimation. **B**: Comparison between the ODE model and the MCell simulations, assuming a homogeneous distribution of the RyRs. **C**: ODE model optimized to fit the experimental data from Segal *et al.* (*22*). Our data is compared to further experimental measurements (*27*) and to other computational models (*19*). **D**: Spine with dendrite used for the ideal case simulations. The mushroom shape spine was extended with a cylinder to simulate the dendrite. **E**: Comparison of the MCell simulations between using only a mushroom shaped spine and using the geometry in A. Mean and standard error of the mean are shown. **F**: Throughout the paper the results will show Ca^2+^ transients at the spine and dendrite. Panel F depicts the geometric compartments used to delineate these measurements.

**Figure S3:**
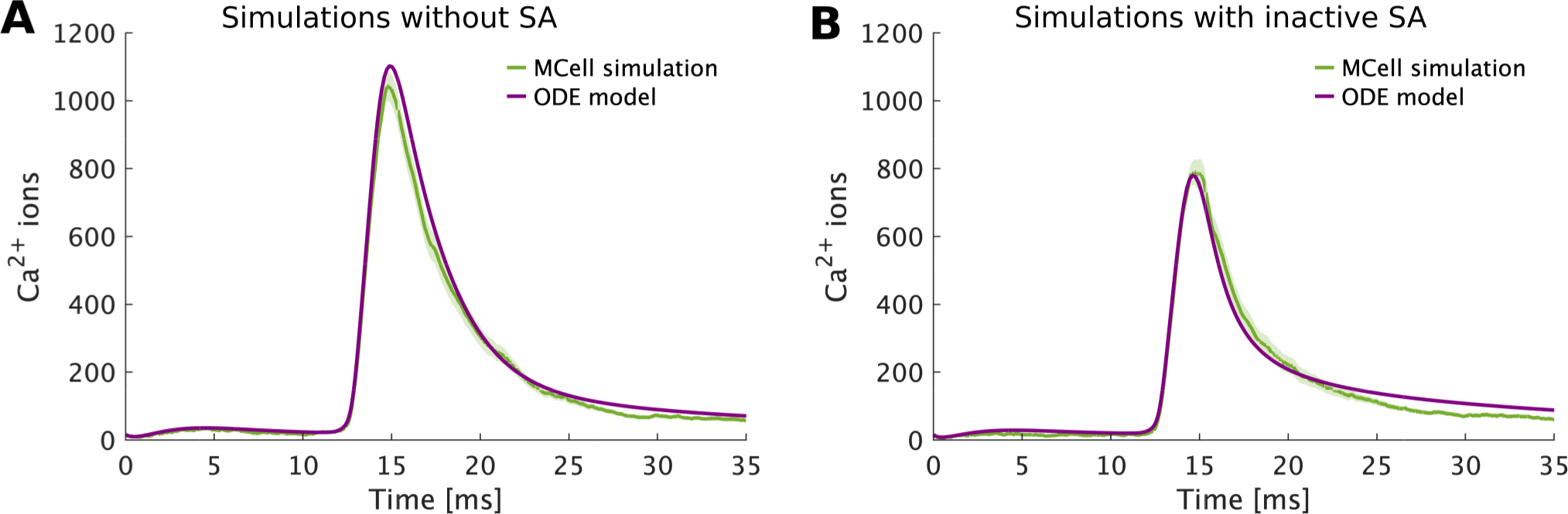
Comparison of ODE model with simulations ran in MCell4. **A**: Simulations were ran without spine apparatus in MCell4 and using the compartmental ODE model. Mean and standard error of the mean are compared. **B**: Simulations were ran with an inactive spine apparatus (only including SERCA) in MCell4 and using the compartmental ODE model. Mean and standard error of the mean are compared.

**Figure S4:**
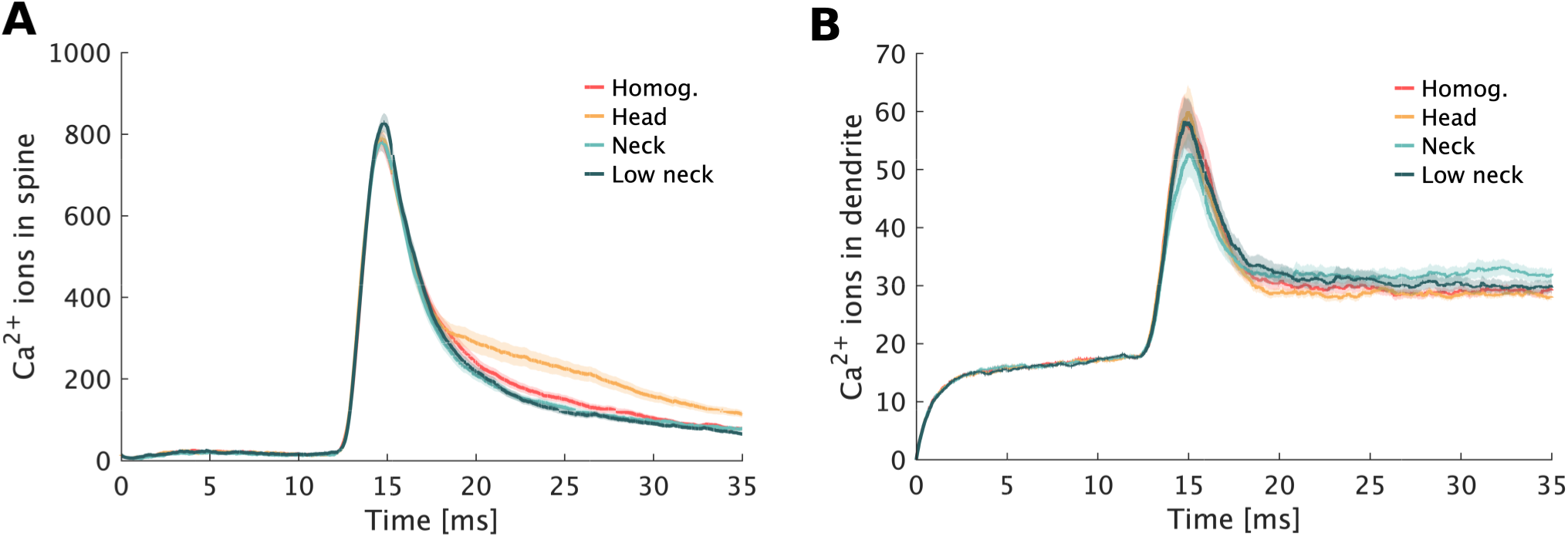
Ca^2+^ dynamics in spine and dendrite averaged over all simulations N for the idealized geometry with idealized ER. **A**: Number of Ca^2+^ ions in the spine averaged over all simulations **N**. **B**: Number of Ca^2+^ ions in the dendrite averaged over all simulations **N**. Mean and standard error of the mean are shown.

**Figure S5:**
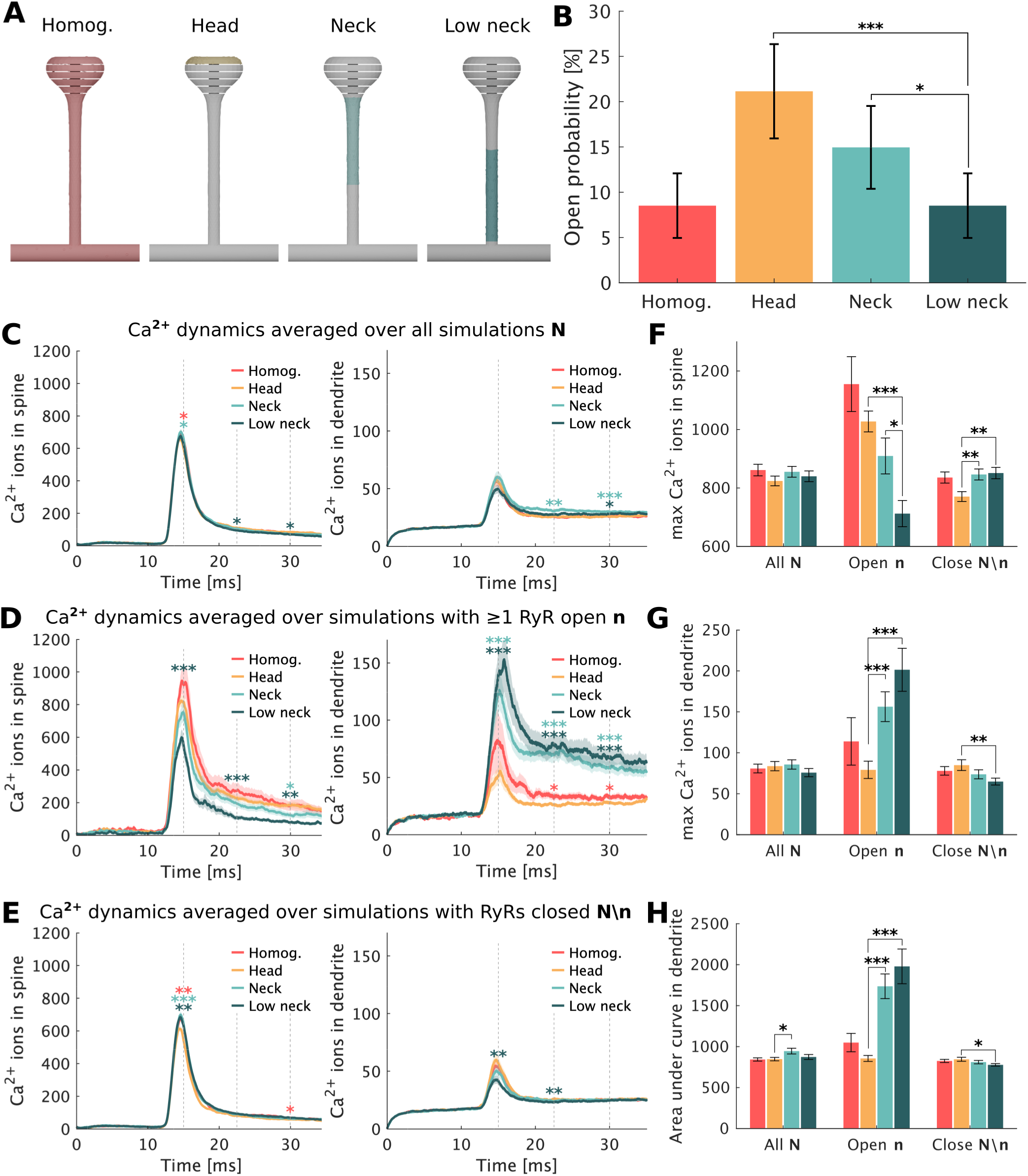
Effect of RyR distribution on ideal spine geometry with an idealized laminar spine apparatus. **A**: RyR distributions considered in the ideal spine with an idealized laminar spine apparatus. **B**: Opening probability for each RyR distribution setup. The bars represent the maximum likelihood estimate of the opening of the RyRs. **C**: Ca^2+^ dynamics in the spine and dendrite averaged over all **N** simulations. Mean and standard error of the mean of all the simulations is shown. **D**: Ca^2+^ dynamics in the spine and dendrite averaged over simulations containing at least one open RyR **n**. Mean and standard error of the mean of all the simulations is shown. **E**: Ca^2+^ dynamics in the spine and dendrite averaged over simulations where all RyRs remained closed **N***\***n**. Mean and standard error of the mean of all the simulations is shown. **F**: Mean of the maximum number of Ca^2+^ ions in the spine. The bars represent the standard error of the mean. **G**: Mean of the maximum number of Ca^2+^ ions in the dendrite. The bars represent the standard error of the mean. **H**: Mean of the area under the curve of Ca^2+^ ions in the dendrite. The bars represent the standard error of the mean.

**Figure S6:**
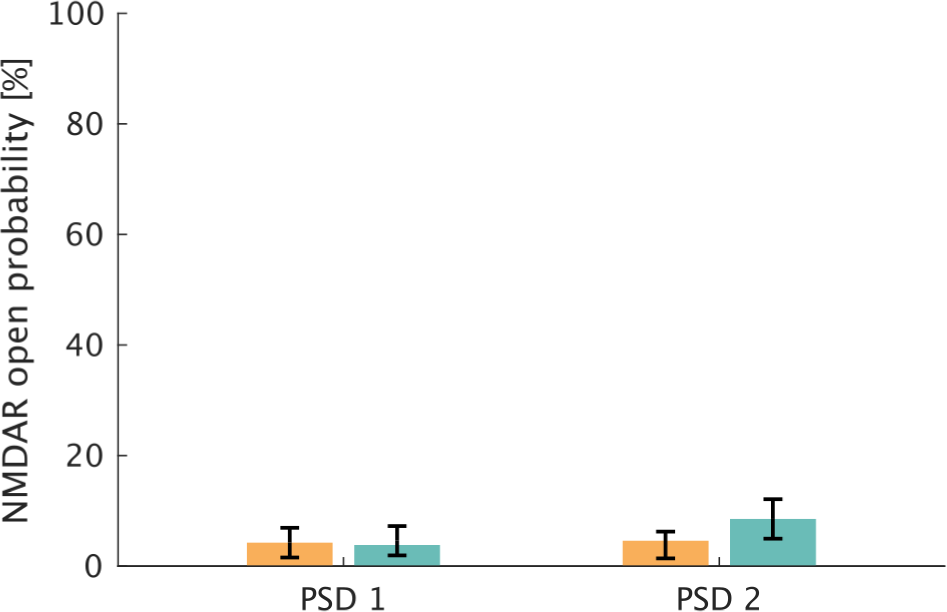
NMDAR opening probability for single PSD stimulation in the realistic plasma membrane geometry.

**Figure S7:**
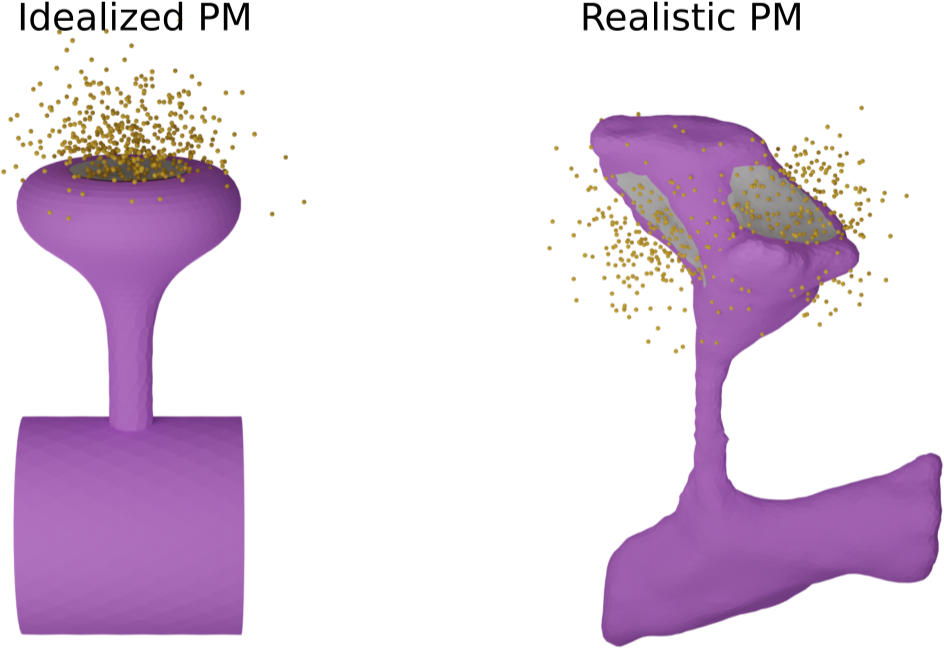
Glutamate diffusion after 0.1 ms in idealized and realistic plasma membrane.

**Figure S8:**
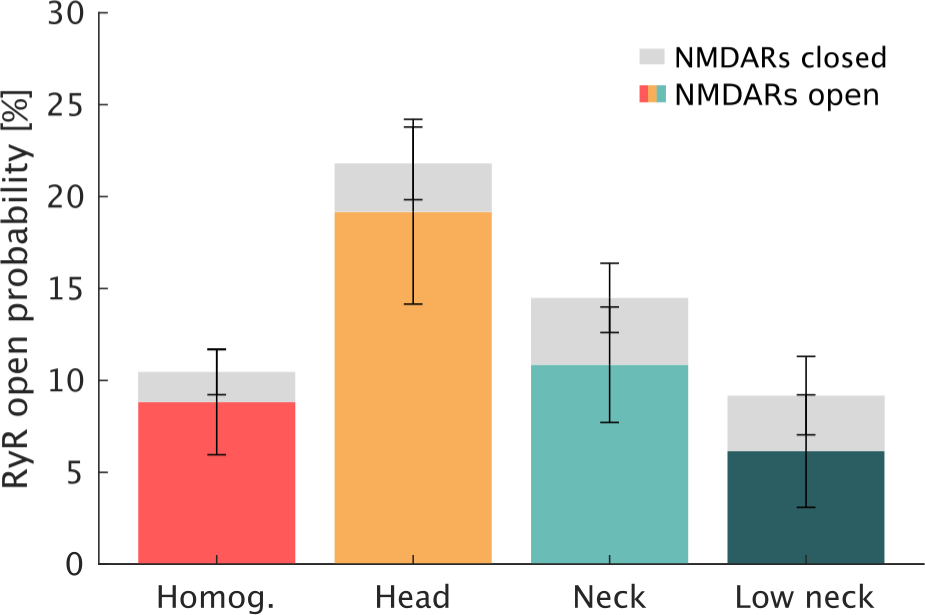
RyR opening probability for each distribution including how many of the simulations contain opened NMDARs and how many closed NMDARs for idealized PM.

**Figure S9:**
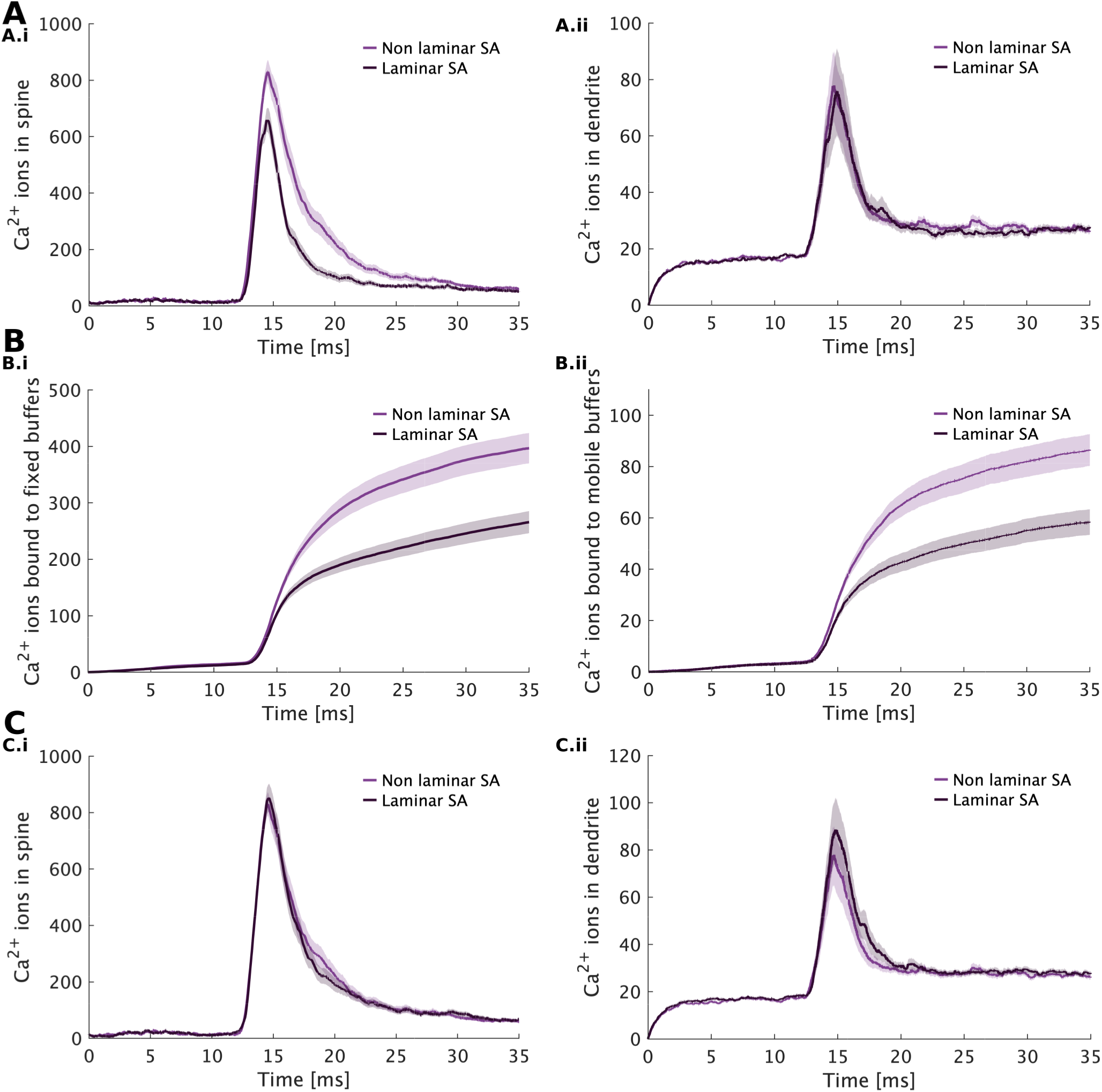
Supplemental comparison of effect of SERCA distribution an Ca^2+^ buffers in spine and dendrite. **A**: Comparison between Ca^2+^ dynamics in an inactive ER (only SERCA) for both laminar and non-laminar ER. **B**: Number of Ca^2+^ ions bound to fixed and mobile buffers in the inactive ER simulation setup. **C**: Ca^2+^ dynamics in an inactive ER (only SERCA) assuming constant number of SERCA.

## References

1 K. M. Harris, F. E. Jensen, B. Tsao, Three-dimensional structure of dendritic spines and synapses in rat hippocampus (CA1) at postnatal day 15 and adult ages: implications for the maturation of synaptic physiology and long-term potentiation. Journal of Neuroscience 12, 2685–2705, DOI 10.1523/JNEUROSCI.12-07-02685.1992 (1992).

2 J. Spacek, K. M. Harris, Three-dimensional organization of smooth endoplasmic reticulum in hippocampal CA1 dendrites and dendritic spines of the immature and mature rat. Journal of Neuroscience 17, 190–203, DOI 10.1523/JNEUROSCI.17-01-00190.1997 (1997).

3 L. A. Colgan, R. Yasuda, Plasticity of dendritic spines: subcompartmentalization of signaling. Annual review of physiology 76, 365–385, DOI 10.1146/annurev-physiol-021113-170400 (2014).

4 M. K. Bell, M. V. Holst, C. T. Lee, P. Rangamani, Dendritic spine morphology regulates calcium-dependent synaptic weight change. Journal of General Physiology 154, DOI 10.1085/jgp.202112980 (2022).

5 E. A. Nimchinsky, B. L. Sabatini, K. Svoboda, Structure and function of dendritic spines. Annual Review of Physiology 64, 313–353, DOI 10.1146/annurev.physiol.64.081501.160008 (2002).

6 P. Rangamani, M. G. Levy, S. Khan, G. Oster, Paradoxical signaling regulates structural plasticity in dendritic spines. Proceedings of the National Academy of Sciences 113, E5298–E5307, DOI 10.1073/pnas.1610391113 (2016).

7 G. Yang, F. Pan, W.-B. Gan, Stably maintained dendritic spines are associated with lifelong memories. Nature 462, 920–924, DOI 10.1038/nature08577 (2009).

8 M. Matsuzaki, N. Honkura, G. C. Ellis-Davies, H. Kasai, Structural basis of long-term potentiation in single dendritic spines. Nature 429, 761–766, DOI 10.1038/nature02617 (2004).

9 A. Van Harreveld, E. Fifkova, Swelling of dendritic spines in the fascia dentata after stimulation of the perforant fibers as a mechanism of post-tetanic potentiation. Experimental Neurology 49, 736–749, DOI 10.1016/0014-4886(75)90055-2 (1975).

10 H. Alimohamadi, M. K. Bell, S. Halpain, P. Rangamani, Mechanical principles governing the shapes of dendritic spines. Frontiers in Physiology 12, 657074, DOI 10.3389/fphys.2021.657074 (2021).

11 M. Bell, T. Bartol, T. Sejnowski, P. Rangamani, Dendritic spine geometry and spine apparatus organization govern the spatiotemporal dynamics of calcium. Journal of General Physiology 151, 1017–1034, DOI 10.1085/jgp.201812261 (2019).

12 D. Ohadi et al., Computational modeling reveals frequency modulation of calcium-cAMP/PKA pathway in dendritic spines. Biophysical Journal 117, 1963–1980, DOI 10.1016/j.bpj.2019.10.003 (2019).

13 M. J. Higley, B. L. Sabatini, Calcium signaling in dendritic spines. Cold Spring Harbor perspectives in biology 4, a005686, DOI 10.1101/cshperspect.a005686 (2012).

14 H. Bading, Nuclear calcium signalling in the regulation of brain function. Nature Reviews Neuroscience 14, 593– 608 (2013).

15 A. K. Lam, A. Galione, The endoplasmic reticulum and junctional membrane communication during calcium signaling. Biochimica et Biophysica Acta (BBA)-Molecular Cell Research 1833, 2542–2559, DOI 10.1016/j.bbamcr.2013.06.004 (2013).

16 P. Jedlicka, A. Vlachos, S. W. Schwarzacher, T. Deller, A role for the spine apparatus in LTP and spatial learning. Behavioural Brain Research 192, 12–19, DOI 10.1016/j.bbr.2008.02.033 (2008).

17 J. N. Bourne, K. M. Harris, Balancing structure and function at hippocampal dendritic spines. Annual Review of Neuroscience 31, 47–67, DOI 10.1146/annurev.neuro.31.060407.125646 (2008).

18 A. Konietzny, S. Wegmann, M. Mikhaylova, The endoplasmic reticulum puts a new spin on synaptic tagging. Trends in Neurosciences, DOI 10.1016/j.tins.2022.10.012 (2022).

19 T. M. Bartol et al., Computational reconstitution of spine calcium transients from individual proteins. Frontiers in Synaptic Neuroscience 7, 17, DOI 10.3389/fnsyn.2015.00017 (2015).

20 Y. Wu et al., Contacts between the endoplasmic reticulum and other membranes in neurons. Proceedings of the National Academy of Sciences 114, E4859–E4867, DOI 10.1073/pnas.1701078114 (2017).

21 A. H. Sharp et al., Differential immunohistochemical localization of inositol 1, 4, 5-trisphosphate-and ryanodine-sensitive Ca2+ release channels in rat brain. Journal of Neuroscience 13, 3051–3063, DOI 10.1523/JNEUROSCI.13-07-03051.1993 (1993).

22 M. Segal, E. Korkotian, Endoplasmic reticulum calcium stores in dendritic spines. Frontiers in Neuroanatomy 8, 64, DOI 10.3389/fnana.2014.00064 (2014).

23 D. F. de Sevilla, A. Núnez, M. Borde, R. Malinow, W. Buno, Cholinergic-mediated IP3-receptor activation induces long-lasting synaptic enhancement in CA1 pyramidal neurons. Journal of Neuroscience 28, 1469–1478, DOI 10.1523/JNEUROSCI.2723-07.2008 (2008).

24 D. F. de Sevilla, W. Buno, The muscarinic long-term enhancement of NMDA and AMPA receptor-mediated transmission at Schaffer collateral synapses develop through different intracellular mechanisms. Journal of Neuroscience 30, 11032–11042, DOI 10.1523/JNEUROSCI.1848-10.2010 (2010).

25 E. Korkotian, M. Segal, Synaptopodin regulates release of calcium from stores in dendritic spines of cultured hippocampal neurons. The Journal of Physiology 589, 5987–5995, DOI 10.1113/jphysiol.2011.217315 (2011).

26 A. Vlachos et al., Synaptopodin regulates plasticity of dendritic spines in hippocampal neurons. Journal of Neuroscience 29, 1017–1033, DOI 10.1523/JNEUROSCI.5528-08.2009 (2009).

27 K. Basnayake et al., Fast calcium transients in dendritic spines driven by extreme statistics. PLoS Biology 17, e2006202, DOI 10.1371/journal.pbio.2006202 (2019).

28 M. Segal, A. Vlachos, E. Korkotian, The spine apparatus, synaptopodin, and dendritic spine plasticity. The Neuroscientist 16, 125–131, DOI 10.1177/1073858409355829 (2010).

29 M. Breit, M. Kessler, M. Stepniewski, A. Vlachos, G. Queisser, Spine-to-dendrite calcium modeling discloses relevance for precise positioning of ryanodine receptor-containing spine endoplasmic reticulum. Scientific Reports 8, 1–17, DOI 10.1038/s41598-018-33343-9 (2018).

30 U. S. Bhalla, Signaling in small subcellular volumes. II. Stochastic and diffusion effects on synaptic network properties. Biophysical Journal 87, 745–753, DOI 10.1529/biophysj.104.040501 (2004).

31 J. H. Kotaleski, K. T. Blackwell, Modelling the molecular mechanisms of synaptic plasticity using systems biology approaches. Nature Reviews Neuroscience 11, 239–251, DOI 10.1038/nrn2807 (2010).

32 T. R. Kolstad et al., Ryanodine receptor dispersion disrupts Ca2+ release in failing cardiac myocytes. Elife 7, e39427, DOI 10.7554/eLife.39427 (2018).

33 X. Shen et al., 3D dSTORM imaging reveals novel detail of ryanodine receptor localization in rat cardiac myocytes. The Journal of Physiology 597, 399–418, DOI 10.1113/JP277360 (2019).

34 M. Herńandez Mesa, J. van den Brink, W. E. Louch, K. J. McCabe, P. Rangamani, Nanoscale organization of ryanodine receptor distribution and phosphorylation pattern determines the dynamics of calcium sparks. PLOS Computational Biology 18, 1–22, DOI 10.1371/journal.pcbi.1010126 (2022).

35 M. Vargas-Caballero, H. P. Robinson, Fast and slow voltage-dependent dynamics of magnesium block in the NMDA receptor: the asymmetric trapping block model. Journal of Neuroscience 24, 6171–6180, DOI 10.1523/JNEUROSCI.1380-04.2004 (2004).

36 P. Jonas, G. Major, B. Sakmann, Quantal components of unitary EPSCs at the mossy fibre synapse on CA3 pyramidal cells of rat hippocampus. The Journal of Physiology 472, 615–663, DOI 10.1113/jphysiol.1993.sp019965 (1993).

37. A. Husar et al., MCell4 with BioNetGen: A Monte Carlo Simulator of Rule-Based Reaction-Diffusion Systems with Python Interface. *bioRxiv*, 2022–05, DOI 10.1101/2022.05.17.492333 (2022).

38. T. M. Inc., MATLAB version: 9.13.0 (R2022b), Natick, Massachusetts, United States, 2022, (https://www.mathworks.com).

39 R. J. Wenthold, R. S. Petralia, A. Niedzielski, et al., Evidence for multiple AMPA receptor complexes in hippocampal CA1/CA2 neurons. Journal of Neuroscience 16, 1982–1989, DOI 10.1523/JNEUROSCI.16-06-01982.1996 (1996).

40 J. C. Magee, D. Johnston, Characterization of single voltage-gated Na+ and Ca2+ channels in apical dendrites of rat CA1 pyramidal neurons. The Journal of Physiology 487, 67–90, DOI 10.1113/jphysiol.1995.sp020862 (1995).

41 A. J. Tanskanen, J. L. Greenstein, A. Chen, S. X. Sun, R. L. Winslow, Protein geometry and placement in the cardiac dyad influence macroscopic properties of calcium-induced calcium release. Biophysical Journal 92, 3379–3396, DOI 10.1529/biophysj.106.089425 (2007).

42 T. Doi, S. Kuroda, T. Michikawa, M. Kawato, Inositol 1, 4, 5-trisphosphate-dependent Ca2+ threshold dynamics detect spike timing in cerebellar Purkinje cells. Journal of Neuroscience 25, 950–961, DOI 10.1523/JNEUROSCI.2727-04.2005 (2005).

43 B. Schwaller, Cytosolic Ca2+ buffers. Cold Spring Harbor Perspectives in Biology 2, a004051, DOI 10.1101/cshperspect.a004051 (2010).

44 P. McPherson, K. Campbell, Characterization of the major brain form of the ryanodine receptor/Ca2+ release channel. Journal of Biological Chemistry 268, 19785–19790, DOI 10.1016/S0021-9258(19)36582-2 (1993).

45 P. S. McPhersonx et al., The brain ryanodine receptor: a caffeine-sensitive calcium release channel. Neuron 7, 17–25, DOI 10.1016/0896-6273(91)90070-G (1991).

46 P. D. Walton et al., Ryanodine and inositol trisphosphate receptors coexist in avian cerebellar Purkinje neurons. The Journal of Cell Biology 113, 1145–1157, DOI 10.1083/jcb.113.5.1145 (1991).

47 M. H. Ellisman et al., Identification and localization of ryanodine binding proteins in the avian central nervous system. Neuron 5, 135–146, DOI 10.1016/0896-6273(90)90304-X (1990).

48 M. D. Stern et al., Local control models of cardiac excitation–contraction coupling: a possible role for allosteric interactions between ryanodine receptors. The Journal of General Physiology 113, 469–489, DOI 10.1085/jgp.113.3.469 (1999).

49 S. Q. Wang, M. D. Stern, E. Ŕıos, H. Cheng, The quantal nature of Ca2+ sparks and in situ operation of the ryanodine receptor array in cardiac cells. Proceedings of the National Academy of Sciences 101, 3979–3984, DOI 10.1073/pnas.0306157101 (2004).

50 N. Singh, T. Bartol, H. Levine, T. Sejnowski, S. Nadkarni, Presynaptic endoplasmic reticulum regulates short-term plasticity in hippocampal synapses. Communications Biology 4, 1–13, DOI 10.1038/s42003-021-01761-7 (2021).

51 Y. Hou et al., Nanoscale organisation of ryanodine receptors and junctophilin-2 in the failing human heart. Frontiers in Physiology 12, 724372, DOI 10.3389/fphys.2021.724372 (2021).

52 D. C. Resasco et al., Virtual Cell: computational tools for modeling in cell biology. Wiley Interdisciplinary Reviews: Systems Biology and Medicine 4, 129–140, DOI 10.1002/wsbm.165 (2012).

53. S. Musall, stdshade, https://www.mathworks.com/matlabcentral/fileexchange/29534-stdshade, [MAT-LAB Central File Exchange; Retrieved August 1], 2023.

54 R. Benavides-Piccione et al., Differential structure of hippocampal CA1 pyramidal neurons in the human and mouse. Cerebral Cortex 30, 730–752, DOI 10.1093/cercor/bhz122 (2020).

55 A. Leung, D. Ohadi, G. Pekkurnaz, P. Rangamani, Systems modeling predicts that mitochondria ER contact sites regulate the postsynaptic energy landscape. NPJ Systems Biology and Applications 7, 26, DOI 10.1038/s41540-021-00185-7 (2021).

56 C. T. Lee et al., 3D mesh processing using GAMer 2 to enable reaction-diffusion simulations in realistic cellular geometries. PLoS Computational Biology 16, e1007756, DOI 10.1371/journal.pcbi.1007756 (2020).

57 W. G. Regehr, D. W. Tank, Postsynaptic NMDA receptor-mediated calcium accumulation in hippocampal CAl pyramidal cell dendrites. Nature 345, 807–810, DOI 10.1038/345807a0 (1990).

58 A. Perez-Alvarez et al., Endoplasmic reticulum visits highly active spines and prevents runaway potentiation of synapses. Nature Communications 11, 5083, DOI 10.1038/s41467-020-18889-5 (2020).

59 M. A. Chirillo, M. S. Waters, L. F. Lindsey, J. N. Bourne, K. M. Harris, Local resources of polyribosomes and SER promote synapse enlargement and spine clustering after long-term potentiation in adult rat hippocampus. Scientific Reports 9, 3861, DOI 10.1038/s41598-019-40520-x (2019).

60 C. Hetz, B. Mollereau, Disturbance of endoplasmic reticulum proteostasis in neurodegenerative diseases. Nature Reviews Neuroscience 15, 233–249, DOI 10.1038/nrn3689 (2014).

61 K. M. Harris, Structural LTP: from synaptogenesis to regulated synapse enlargement and clustering. Current Opinion in Neurobiology 63, 189–197, DOI 10.1016/j.conb.2020.04.009 (2020).

62 R. Yasuda, B. L. Sabatini, K. Svoboda, Plasticity of calcium channels in dendritic spines. Nature Neuroscience 6, 948–955, DOI 10.1038/nn1112 (2003).

63 J. C. Magee, D. Johnston, A synaptically controlled, associative signal for Hebbian plasticity in hippocampal neurons. Science 275, 209–213, DOI 10.1126/science.275.5297.209 (1997).

64 R. Yuste, W. Denk, Dendritic spines as basic functional units of neuronal integration. Nature 375, 682–684, DOI 10.1038/375682a0 (1995).

65 H. Falahati, Y. Wu, V. Feuerer, H.-G. Simon, P. De Camilli, Proximity proteomics of synaptopodin provides insight into the molecular composition of the spine apparatus of dendritic spines. Proceedings of the National Academy of Sciences 119, e2203750119, DOI 10.1073/pnas.2203750119 (2022).

66 K. Obashi, J. W. Taraska, S. Okabe, The role of molecular diffusion within dendritic spines in synaptic function. Journal of General Physiology 153, e202012814, DOI 10.1085/jgp.202012814 (2021).

67 Y. Wang, J. Wu, M. J. Rowan, R. Anwyl, Ryanodine produces a low frequency stimulation-induced NMDA receptor-independent long-term potentiation in the rat dentate gyrus in vitro. The Journal of Physiology 495, 755–767, DOI 10.1113/jphysiol.1996.sp021631 (1996).

68 C. R. Raymond, S. J. Redman, Different calcium sources are narrowly tuned to the induction of different forms of LTP. Journal of Neurophysiology 88, 249–255, DOI 10.1152/jn.2002.88.1.249 (2002).

69 I. Valdés-Undurraga et al., Long-term potentiation and spatial memory training stimulate the hippocampal expression of RyR2 calcium release channels. Frontiers in Cellular Neuroscience 17, 1132121, DOI 10.3389/fncel.2023.1132121 (2023).

70 I. Vega-Vásquez et al., Hippocampal dendritic spines express the RyR3 but not the RyR2 ryanodine receptor isoform. Biochemical and Biophysical Research Communications 633, 96–103, DOI 10.1016/j.bbrc.2022.10.024 (2022).

71 N. Emptage, T. V. Bliss, A. Fine, Single synaptic events evoke NMDA receptor–mediated release of calcium from internal stores in hippocampal dendritic spines. Neuron 22, 115–124, DOI 10.1016/s0896-6273(00)80683-2 (1999).

72 M. K. Park, O. H. Petersen, A. V. Tepikin, The endoplasmic reticulum as one continuous Ca2+ pool: visualization of rapid Ca2+ movements and equilibration. The EMBO journal 19, 5729–5739, DOI 10.1093/emboj/19.21.5729 (2000).

73 J. A. Wasserstrom et al., Variability in timing of spontaneous calcium release in the intact rat heart is determined by the time course of sarcoplasmic reticulum calcium load. Circulation Research 107, 1117–1126, DOI 10.1161/CIRCRESAHA.110.229294 (2010).

74 P. J. Dittmer, A. R. Wild, M. L. Dell’Acqua, W. A. Sather, STIM1 Ca2+ sensor control of L-type Ca2+-channel-dependent dendritic spine structural plasticity and nuclear signaling. Cell Reports 19, 321–334, DOI 10.1016/j.celrep.2017.03.056 (2017).

75 T. Mäki-Marttunen, N. Iannella, A. G. Edwards, G. T. Einevoll, K. T. Blackwell, A unified computational model for cortical post-synaptic plasticity. Elife 9, e55714, DOI 10.7554/eLife.55714 (2020).

76 M. K. Bell, P. Rangamani, Crosstalk between biochemical signalling network architecture and trafficking governs AMPAR dynamics in synaptic plasticity. The Journal of Physiology, DOI 10.1113/JP284029 (2023).

77 G. Zhu, Y. Liu, Y. Wang, X. Bi, M. Baudry, Different patterns of electrical activity lead to long-term potentiation by activating different intracellular pathways. Journal of Neuroscience 35, 621–633, DOI 10.1523/JNEUROSCI.2193-14.2015 (2015).

78 T. M. Dhanrajan et al., Expression of long-term potentiation in aged rats involves perforated synapses but dendritic spine branching results from high-frequency stimulation alone. Hippocampus 14, 255–264, DOI 10.1002/hipo.10172 (2004).

79 P. J. Dittmer, M. L. Dell’Acqua, W. A. Sather, Synaptic crosstalk conferred by a zone of differentially regulated Ca2+ signaling in the dendritic shaft adjoining a potentiated spine. Proceedings of the National Academy of Sciences 116, 13611–13620, DOI 10.1073/pnas.1902461116 (2019).

